# Mechanistic Insights into Unfolding of the Mitochondrial Localization Sequence M1 of TDP-43 and Identification of a MAPKAPK2 Inhibitor as a potential binder of M1 for inhibition of TDP-43’s Aberrant Mitochondrial Localization

**DOI:** 10.64898/2026.06.17.732848

**Authors:** Ramkumar Balaji, Shanu Bhardwaj, Jessica Baa, Himanshu Joshi, Basant K. Patel

## Abstract

Mechanistic elucidation and inhibition of the pathogenic aberrant mitochondrial localization of the RNA/DNA-binding protein, TDP-43, can help in the therapeutics of the neurodegenerative disease amyotrophic lateral sclerosis (ALS). A mitochondrial localization sequence of TDP-43, M1, is largely solvent inaccessible, therefore, how it interacts with the mitochondrial import machinery to facilitate TDP-43’s transit to mitochondria is unclear. Towards this, we examined the unfolding TDP-43’s N-terminal domain (NTD) that hosts M1, using equilibrium all-atom molecular dynamics (MD) simulations, and observed an early loss of the hydrogen-bonded interactions between β4-β5 bridge and the interactions involving residues Phe-35 and Gly-40 of M1, indicating structural lability of M1 to become solvent-accessible that may enhance its interaction with the mitochondrial receptor(s) for import. Furthermore, *via* virtual screening of 2,115 FDA-approved and 515,545 non-FDA-approved small molecules from ZINC15 database towards binding to M1 and inhibiting TDP-43’s mitochondrial import, we identified a molecule, ZINC73240059, that was previously characterized as an inhibitor of MAP kinase-activating protein kinase 2 (MAPKAPK2). ZINC73240059 remains stably bound to M1 of NTD during MD simulations manifesting negative Gibbs free energy (ΔG) with significant contribution from Pro-36 of M1. Overall, ZINC73240059 can be a molecule of interest towards thwarting TDP-43’s pathogenic mitochondrial localization in ALS.

## 1.0 Introduction

Mislocalization to cytoplasm and aggregation of the nuclear-resident TAR DNA-binding protein (TDP-43) are associated with several neurodegenerative diseases, including amyotrophic lateral sclerosis (ALS) and frontotemporal lobar degeneration (FTLD) and certain other neurodegenerative diseases. ^1,2^ TDP-43 is a 414 amino acid DNA/RNA-binding protein comprising a folded N-terminal domain (NTD), conserved bi-partite RNA recognition motif (RRM1-2), and an aggregation-prone C-terminal domain (CTD) harboring a prion-like glutamine/asparagine-rich sequence. ^3^ Mutations in TDP-43 are linked to both sporadic and familial ALS. Additionally, the C-terminal fragments of post-translationally modified TDP-43 are reported to seed the pathological aggregation of TDP-43 in the neuronal and glial cells of the affected ALS patients. ^4–6^ Several studies have investigated the mechanisms and targeting of the pathological aggregation of TDP-43 and its C-terminal fragments. ^7–12^

Apart from a nuclear localization signal (NLS) and a nuclear export signal (NES), TDP-43 sequence was predicted to also harbor six internal potential mitochondrial localization signal sequences (M1-M6) of which three sequences, M1, M3, and M5 were found to facilitate the aberrant localization of TDP-43 into mitochondria. ^13^ In fact, in addition to the cytoplasmic mislocalization and aggregation, TDP-43 was also reported to mislocalize to mitochondria of neurons of the ALS/FTLD patients ^13^, leading to mitochondrial dysfunction, as evidenced by decreased mitochondrial membrane potential ^14^, increased oxidative stress, ^15–18^ suppressed mitochondrial complex I activity, and reduced mitochondrial ATP synthesis. ^19^ Although various reports suggest TDP-43-induced mitochondria-mediated toxicity, extensive studies examining the mechanisms by which mitochondrial signals are exposed to mediate the TDP-43 internalization into mitochondria, are lacking. A recent study showed that overexpression of the mitophagy receptor, FUNDC1 enhanced the TDP-43-TOM70 and TOM70-DNAJA2(HSP40) interactions, suggesting that this interaction promotes the recognition and import of TDP-43/HSPA8(Hsp70)/DNAJA2(Hsp40) by the TOM70 channel. ^20^ Transmembrane proteins of the inner membrane contain internal mitochondrial localization sequence and undergo partial unfolding before import into the mitochondria through the TOM70-TIM22 complex channels in the outer and inner mitochondrial membrane respectively. ^21^ Although, TDP-43 does not belong to the transmembrane class of proteins, the presence of internal mitochondrial sequences and TDP-43’s reported interaction with TIM22 ^22^ suggest a possibility of chaperone-mediated unfolding event to facilitate its mitochondrial import.

Earlier molecular dynamics (MD) simulation studies on the folding and unfolding of NTD of TDP-43 at higher temperatures and in the presence of 8 M urea or 8 M DMSO illustrated partially unfolded states and their role in dimerization/oligomerization of the protein. ^23,24^ An experimental study using the intrinsic fluorescence of a core amino acid residue, Trp-68, as a probe also confirmed the presence of an intermediate state formed before unfolding at 4.5 M urea in TDP-43. The intermediate state was reported to form even in the monomeric state which remained unaffected by the oligomerization status. ^25^ However, the effect of unfolding on the conformation of TDP-43’s internal mitochondrial signal sequences remains to be investigated.

Highlighting the importance of the mitochondrial damage by TDP-43 to cause toxicity, deletions of a mitochondrial fission protein (Dnm1) and the mitochondrial pro-apoptotic protein (Ybh3) rescued the TDP-43 toxicity, oxidative stress and cell death in a yeast heterologous expression model. ^26^ In addition, certain small molecules, including antioxidants such as N-acetylcysteine (NAC), curcumin, and ascorbic acid, are effective in alleviating oxidative stress-mediated TDP-43 toxicity in various neurodegenerative conditions. ^27^ Strikingly, inhibition of the aberrant mislocalization of TDP-43 *via* utilizing TAT-fused peptide inhibitors mimicking either M1 (aa: 35-41, FPCAGL) or M3 (aa: 146-150, GFGFV) internal mitochondrial signals of TDP-43 could alleviate the neuronal loss in transgenic mice expressing TDP-43 possibly by competitively binding to the mitochondrial membrane import complexes. ^13^ Thus, targeting TDP-43’s internal mitochondrial localization signal sequences with small molecules that could stably bind to these sequences and prevent TDP-43’s recognition by the mitochondrial import receptors, appears as a potential therapeutic approach against neurodegeneration induced by TDP-43 proteinopathy. ^13^ In fact, Vitamin D3 was recently found *in silico* to stably interact with M3 mitochondrial localization sequence of TDP-43. ^28^

Structure-based virtual drug screening and MD simulation are widely employed powerful computational techniques for modeling the interactions among molecules or the atoms within a molecule. These techniques reduce the search space for conventional laboratory-based screening of small-molecule inhibitors to biomolecules. Structure-based virtual screening ranks molecules being tested for their binding affinity to a known target macromolecule based on docking scores. ^29^ MD simulations can sample the conformations of biomolecules as they fold or unfold under different environmental conditions, and as they interact with other molecules. ^30^ All-atom MD simulations are employed to understand the protein’s dynamic motion, which cannot be investigated at the atomic level by experimental techniques that provide static structural information. ^31^ MD simulations are also gaining importance for investigating the aggregation pathways of proteins that form amyloids and the phase separation behavior of proteins involved in neurodegeneration. ^32–34^

In this study, to gain mechanistic insights into how a normally solvent inaccessible mitochondrial localization sequence, M1, facilitates TDP-43’s mitochondrial import, we performed equilibrium all-atom MD simulations on the N-terminal domain of TDP-43 at 310 K, 350 K and 450 K in presence of 4 M urea to accelerate conformational sampling and study its unfolding with emphasis on M1. This enabled characterization of the time-dependent structural perturbations in the M1 region as the protein unfolds. The structure of the full-length TDP-43 is not yet determined experimentally due to the poor solubility and the presence of a long aggregation-prone intrinsically disordered C-terminal region in TDP-43. However, structures of the NTD and RRM1-2 domains of TDP-43 deciphered using X-ray crystallography, nuclear magnetic resonance (NMR) spectroscopy or cryo-electron microscopy (Cryo-EM), are available. We utilized the available NMR solution structure (PDB ID: 2N4P ^35^, aa: 1-77) and X-ray crystallographic structures (PDB IDs: 5MDI ^36^ and 6T4B ^37^, aa: 2-79) of NTD in this study. Furthermore, we performed a large-scale virtual screening of 2,115 FDA-approved and 515,545 non-FDA-approved small molecules from the ZINC15 database ^38^ towards drug-repurposing to target and inhibit the M1-mediated mitochondrial import of TDP-43 *via* finding small molecules that can stably bind to M1. The stability of the complexes formed by the screened-out binders of M1 was further evaluated by all-atom MD simulations in explicit water solvent at 310 K followed by structural and thermodynamic stability analyses (**Figure 1**).

**Figure 1:**
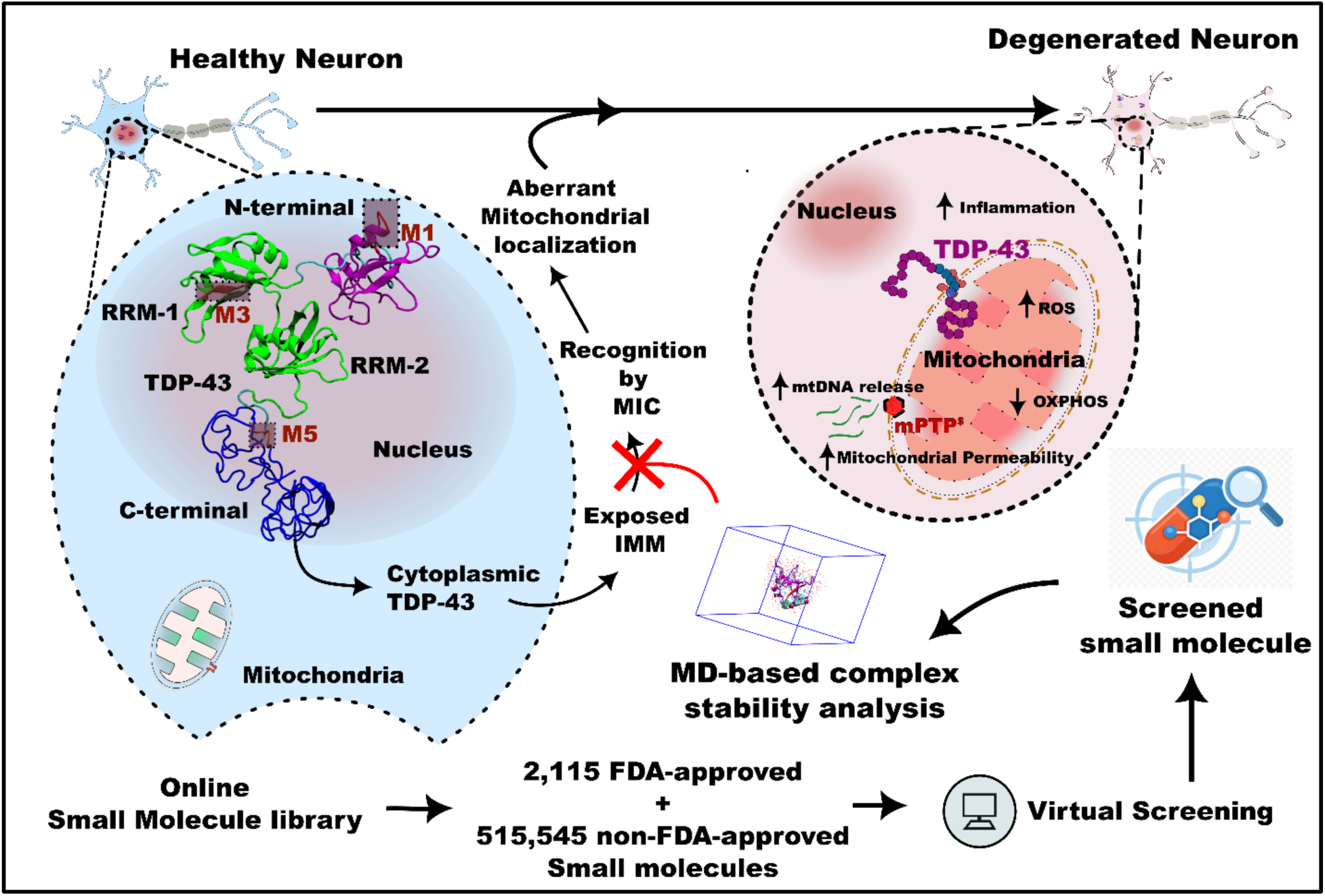
Schematic representation of *in silico* approach to screen and validate small molecules with potential of inhibiting the mitochondrial localization of TDP-43 to mitigate the induced neurotoxicity. Healthy neurons have the RNA/DNA-binding protein, TDP-43, in the nucleus under homeostatic cellular conditions with intact mitochondria. However, cytoplasmic mislocalization could expose the internal mitochondrial signal sequences (IMM), M1, M3, and M5 to recognition by mitochondrial import complexes (MIC), leading to the protein’s aberrant mitochondrial localization. This aberrant mitochondrial mislocalization of TDP-43 has been reported in neurodegenerative diseases, leading to perturbation in mitochondrial dynamics and function. These perturbations include increased reactive oxygen species (ROS), decreased ATP production by oxidative phosphorylation (OXPHOS), increased mitochondrial permeability, and, hence, distorted mitochondrial membrane potential, mitochondrial DNA (mtDNA) release into the cytoplasm through the mitochondrial permeability transition pore (mPTP) complex, leading to inflammation and apoptotic signaling. In this study, large-scale virtual screening of small molecules from an online library is performed to target the M1 region. All-atom MD simulation is employed to further evaluate the stability of the complex between the N-terminal domain containing the M1 sequence and the screened molecules. A small molecule that is identified to stably interact with M1 of TDP-43 can be proposed as an inhibitor of the mitochondrial localization of TDP-43 that could potentially rescue neuronal degeneration *via* blocking the TDP-43’s aberrant mitochondrial mislocalization.

## 2.0 Results and Discussion

### Unfolding of TDP-43 N-terminal domain reveals the hydrogen-bonded network involving the residues of internal mitochondrial localization sequence, M1, is labile

In the native structure, the mitochondrial localization sequence, M1 (comprising of: Phe-35, Pro-36, Gly-37, Ala-38, Cys-39, Gly-40, Leu-41), present in the NTD of TDP-43 is partially buried with three of its residues completely solvent inaccessible,^35^ which is counterintuitive to its ability to interact with the mitochondrial receptors for facilitating the import of TDP-43. Possibly, NTD may undergo partial unfolding assisted by chaperones in order to expose M1 to solvent for this purpose. Therefore to gain insights, we examined the unfolding events of NTD, with focus on M1, using its three available starting conformations from the RCSB PDB database: the solution NMR structure (PDB ID: 2N4P, aa: 1-77) after the removal of the 6x histidine tag ^21^ and two X-ray crystallographic structures (PDB ID: 6T4B,^37^ aa: 2-79 and 5MDI,^36^ aa: 2-79). When examined for their structural similarity, the solution NMR structure, 2N4P, showed a backbone root-mean-squared deviation (RMSD) of about 3.8 Å from the two crystal structures (6T4B and 5MDI), while both crystal structures were quite superimposable manifesting very minimal differences (**Supplementary Figure SF1a-b)**. However, the overall percentage relative solvent accessibilities (%RSA) of the amino acid residues were similar in these three structures and in fact, the major secondary structural elements and motifs were also found preserved (**Supplementary Figure SF1c**).

To document the unfolding events of NTD, we solvated the NMR structure (2N4P) and the crystal structure (6T4B) in explicit water, in absence and presence of 4 M urea. Then, equilibrium molecular dynamics (MD) simulations were carried out at 310, 350, and 450 K to visualize the unfolding of NTD at the atomic level using the systems summarized in **Table 1** and for visualization, one of the fully assembled, simulated system containing 2N4P structure in explicit urea solution is depicted in **Figures 2a** and **2b**.

**Figure 2:**
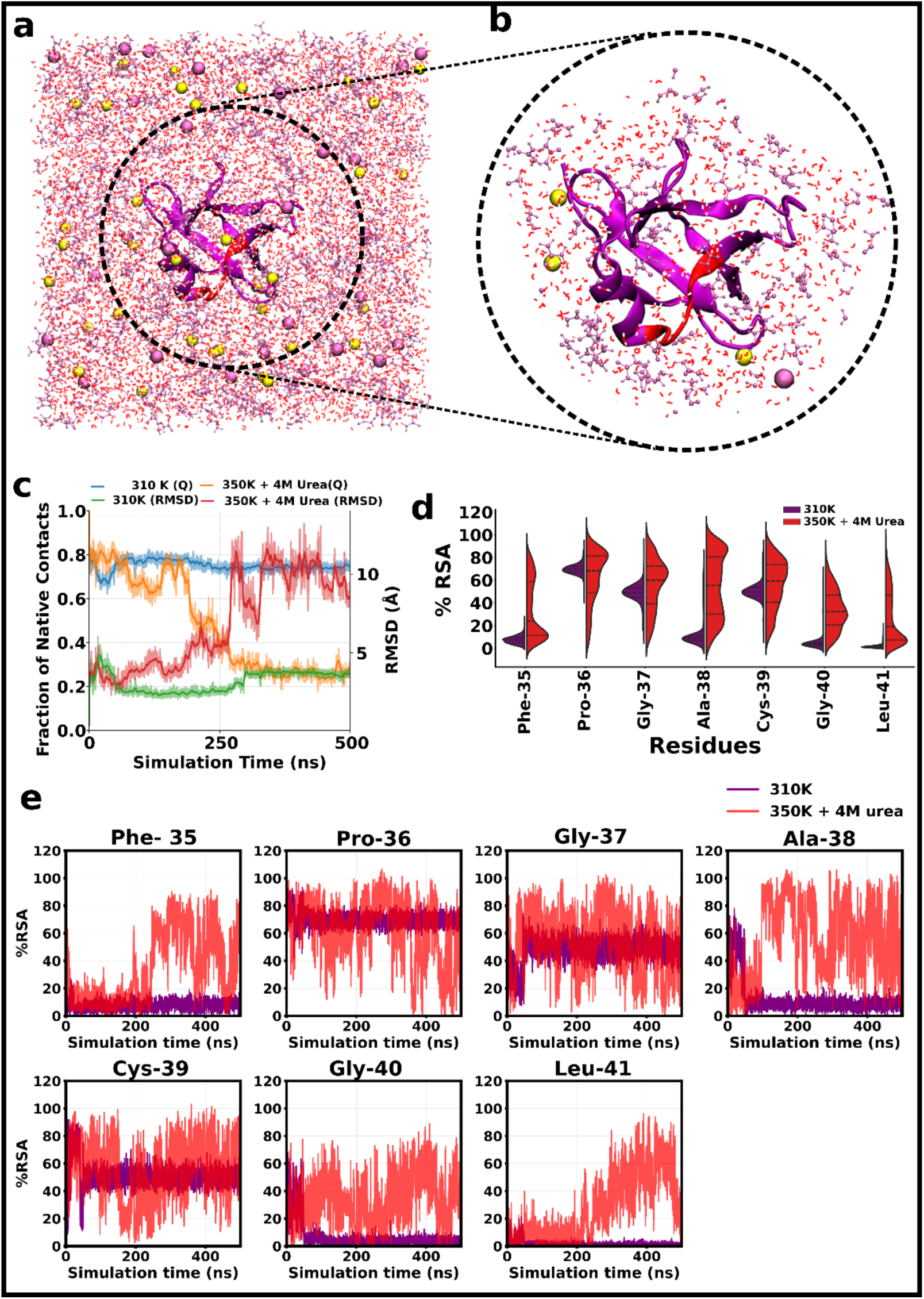
MD simulation analysis of the N-terminal domain of TDP-43 (PDB ID: 2N4P) at 310 K and 350 K+4 M urea simulation conditions. **a)** An instantaneous snapshot of the all-atom MD simulation model, showing the NTD of TDP 43 protein (2N4P) in purple with M1 residues in red, denaturant urea in mauve ball-stick representation. Na^+^ and Cl^−^ions are shown in yellow and mauve sphere representations, respectively, while water is shown in red and white lines. **b)** Zoomed-in representation of the protein and surrounding water molecules. **c)** The fraction of native contacts (Q) (left Y-axis) and RMSD profile (right Y-axis) of the 2N4P at 310 K and 350 K + 4 M urea conditions as a function of simulation time, shown in a twin Y axis plot. **d)** Split violin plot showing the fluctuations in the percentage relative solvent accessibility of the residues of the M1 region at 310 K and 350 K + 4 M urea simulation conditions. **e)** Time evolution of the percentage relative solvent accessibility (%RSA) of the residues of the M1 region in the 310K and 350K + 4M urea simulations.

**Table 1:** Molecular dynamics simulation systems for TDP-43 NTD used in this study.

| TDP-43 NTD Protein Structure (PDB ID) | No. of urea Molecules | Number of atoms in the system* | Temperature (K) |
| --- | --- | --- | --- |
| 2N4P | 0 | 21,571 | 310 |
|  | 760 | 29,410 | 350 |
|  | 0 | 30,764<sup>#</sup> | 450 |
|  | 760 | 29,434<sup>#</sup> | 450 |
| 6T4B | 0 | 21,403<sup>#</sup> | 310 |
|  | 750 | 29,183<sup>#</sup> | 350 |
|  | 760 | 29,284<sup>#</sup> | 400 |
|  | 760 | 29,284<sup>#</sup> | 450 |

\*Number of atoms in the system = No. of protein atoms + No. of water solvent atoms + No. of ion atoms + No. of urea atoms.
<sup>#</sup>Simulation results are available in the supplementary document of the manuscript.

Earlier experimental studies on the NTD of TDP-43 reported unfolding intermediate states at urea concentrations as low as 4.5 M, whereas the protein domain could maintain its fold when incubated at temperatures from 293 to 315 K ^25^. To characterize the unfolding of the NTD of TDP-43, we performed MD simulations at high temperature and in the presence of 4 M urea. In our simulations, the 2N4P structure unfolded at 350 K in the presence of 4 M urea (**Figure 2c**). A transition from folded to unfolded state, elucidating the crucial intermediates, can be visualized in **<u>Supplementary</u> <u>Movie S1</u>**. When the time evolution of the fraction of native contacts (Q value) and the root-mean-square deviation (RMSD) were used to track conformational changes during unfolding (**Figure 2c**), the Q value for the 310 K equilibrium simulations of the 2N4P structure stabilized around 0.7, while the RMSD stabilized around 4 Å. The simulation at 350 K + 4 M urea for 2N4P showed a decline in the Q value after ∼190 ns, followed by stabilization at 0.2 after ∼280 ns. Interestingly, the decrease in the Q value was gradual, illustrating the snapshots of the unfolding intermediates. During the simulation time interval of 210-260 ns, the Q value exhibited a brief steady phase around 0.5, indicating an intermediate state. A similar trend was observed in the increase in RMSD for the 2N4P structure during the 350 K + 4 M urea simulation. The RMSD fluctuated until 210 ns, after which a steady phase at approximately 5 Å was observed from 210 ns to 260 ns, followed by significant fluctuations until the end of the simulation. Lower native contact and higher RMSD quantify the unfolding of 2N4P at 350 K in the presence of 4 M urea, which is also evident from the visualization of the MD simulation trajectories **(<u>Supplementary Movie S1</u>).** The MD simulations at higher temperature with or without urea had shown instantaneous complete unfolding without intermediate states accumulation and hence were not further analyzed **(Supplementary Figure SF2)**.

When the relative solvent accessibility (%RSA) was examined to analyze the fluctuations in the M1 residues upon unfolding, the probability distribution of the %RSA indicated that the residues of M1, which were buried during the 310 K equilibrium MD simulation of the 2N4P structure, began to expose to solvent during the 350 K + 4 M urea MD simulation (**Figure 2d, e**). Among these, Gly-40 fluctuated from 50 ns, followed by Ala-38, Phe-35, and Leu-41 at approximately 100 ns, 220 ns, and 250 ns, respectively, indicating that they were affected at the early unfolding events. The minimum distances of hydrogen-bonded amino acid pairs as a function of time were compared for the folded (310 K) and unfolding (350 K with 4 M urea) simulations to better understand the conformational dynamics of the N-terminal domain during the transition. The box plot of the distances between hydrogen-bonded residue pairs is shown at 50 ns intervals in the 500 ns folded and unfolding simulations (**Supplementary Figure SF3**). The increased interquartile range (IQR) in the 350 K + 4 M simulations relative to the 310 K simulations within a particular time interval was interpreted as a fluctuation in hydrogen-bond interactions between residue pairs, suggesting that these interactions were affected at unfolding conditions. Fluctuations in the minimum pairwise distances, averaged over 50 ns intervals, indicate that interactions involving the terminal residues of the α1 helix–encompassing part of the M1 region as well as the β4 (aa: 55–57)–β5 (aa: 60–62) strands and adjacent loop regions, are disrupted at early stages of unfolding (**Figure 3, Supplementary Figure SF3** and **<u>Supplementary Movie S1</u>**). The fluctuations in the β4-β5 bridge interactions were accompanied by increased movement of the β4 strand, leading to the greater instability in the α1-loop-β3 region, which includes the M1 residues. This further affected hydrogen-bonding interactions among the residues involved in the α1-helix, indicating unfolding. In addition to these residue pairs in the secondary structures and in the loop regions, amino acids that are far away in the sequence of the N-terminal region were also reported to form hydrogen-bonded pairs. These include Leu28-Gly59, Val26-Leu61, Cys39-Asn76, Tyr43-Arg52, Val54-Leu41, Arg6-Gly69, Leu71-Val7, Tyr73-Thr8, Val75-Glu9, and these hydrogen-bonded pairs did not show much fluctuation in their pair-distances until 250 ns of the simulation time. As also reported previously ^35^, the salt bridge interaction between residues Asp-23 and Arg-52 was only disrupted after 250 ns of the simulation time (**Supplementary Figure SF3**).

**Figure 3:**
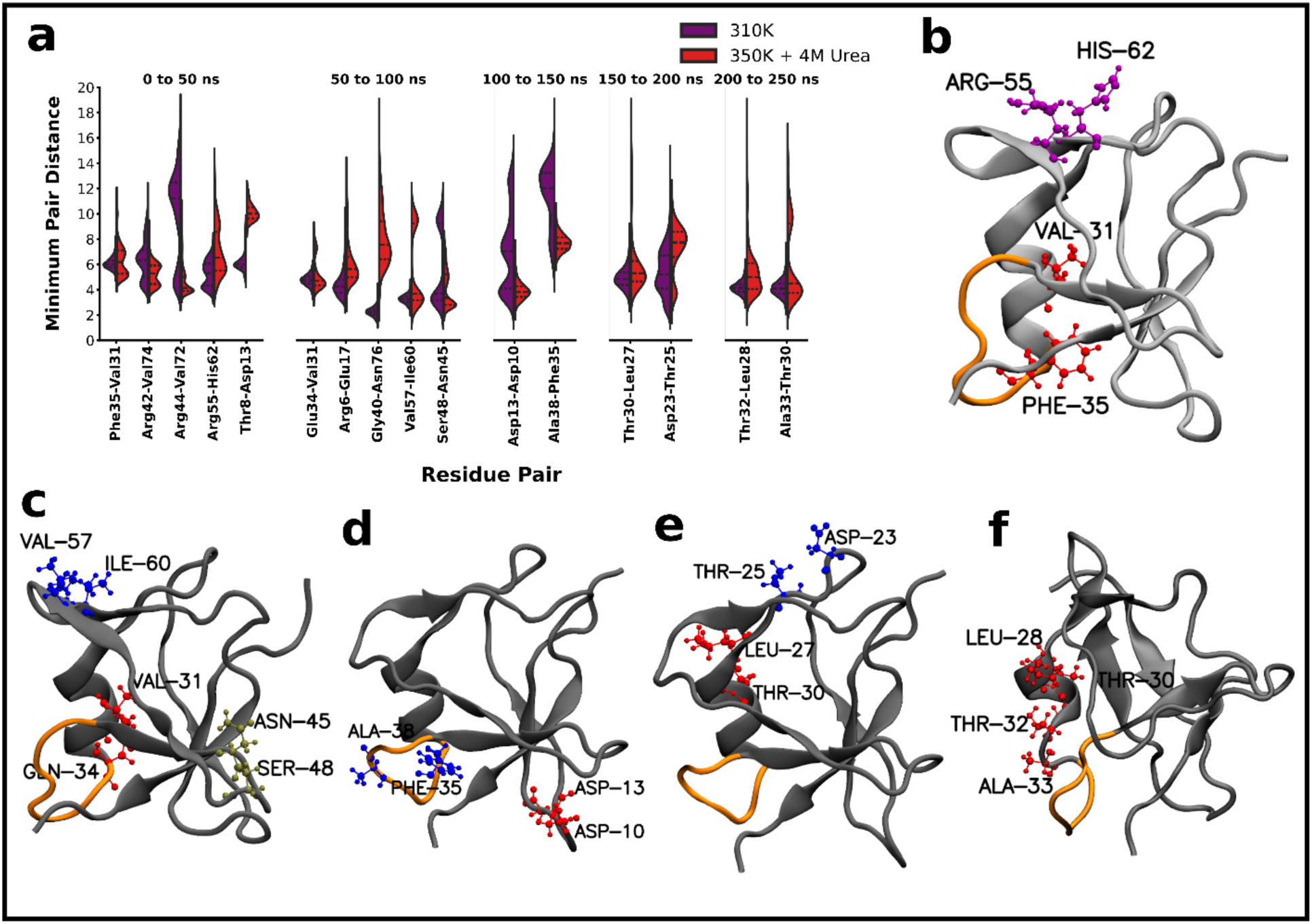
Pair distances between the residues in hydrogen bonding in the N-terminal domain structure of TDP-43 (PDB ID: 2N4P) throughout the MD simulation time. **a)** The distribution of the minimum distances between the residue pairs that showed large variance during the simulation of the N-terminal TDP-43 averaged for 0-50, 50-100, 150-200, 200-250 ns windows during the simulation time, shown as a split violin plot for 310 K (purple) and 350 K + 4M urea (red). **b-f)** Instantaneous snapshots of the protein during the simulation highlighting the amino-acid pairs that showed greater fluctuations at 350 K in 4 M urea simulations are shown in different colors. The New cartoon representation of the protein is shown in cyan, with the M1 residues marked in red. The simulation snapshots represented for the protein are from 0-50, 50-100, 100-150, 150-200, 200-250 ns, respectively. The simulation beyond this showed fluctuations in almost all the hydrogen-bonded residue pairs indicating unfolding.

At the onset of unfolding, the %RSA of Gly-40, Ala-38, and Phe-35 may reflect that hydrogen-bonded interactions involving the M1 residues are affected. In contrast, the residue Leu-41, which is involved in a hydrogen-bonded interaction with Val-54, became solvent-accessible after only 250 ns of simulation time. The residues Phe-35, Ala-38, Gly-40, and Leu-41 are among the residues that are reported to form the hydrophobic core of the N-terminal ^35,39^, which are also required to maintain the N-terminal’s stability. The observations here, hence, suggest that the residues Phe-35, Ala-38, and Gly-40 may be driving the unfolding events through intermediate states. At the same time, Leu-41 shows fluctuations that appear critical for unfolding but not for intermediate-state formation. The gromos clustering, performed every 50 ns along the trajectories, further confirmed the sequence of events in the unfolding of this protein domain. (**Figure 3** and **<u>Supplementary Movie S1</u>**).

In contrast, the 6T4B structure’s MD simulation at 450 K in the presence of 4 M urea did not show any intermediate states but began unfolding within 30 ns **(Supplementary Figure SF4a).** No unfolding was observed at temperatures below 450K, even in the presence of 4M urea **(Supplementary Figure SF4b-d)**. The Q value steadily declined to below 0.1 and stabilized after 30 ns, while the RMSD began to fluctuate, reaching a maximum of 40Å (**Supplementary Figure SF4a**). Moreover, the %RSA reflected the same trend, with all residues of M1 fluctuating and becoming solvent-exposed from above ∼30 ns (**Supplementary Figure SF4e, f**). Hence, the pair distances of the residues reported in hydrogen bonding were measured for the trajectories at intervals of 5ns until 25 ns, after which the entire remaining trajectory was used (**Supplementary Figure SF5a-h**). The pair distances calculated for 6T4B unfolding conditions also showed a similar trend to that observed in the 2N4P structure.

Based on the analysis of the MD simulation trajectories, we propose that perturbations in hydrogen-bonded interactions involving the residues of the M1 and those among the β4-β5 bridge initiate unfolding of the NTD of TDP-43. The loss of interactions observed in the α1-loop region comprising the M1 residues can be attributed to the α-helix’s ability to undergo local conformational changes when near the native state. However, the interactions within the β-sheet are relatively stable and are disrupted only concomitantly with global unfolding and tertiary-structural changes. ^40^

### Virtual screening by docking identifies FDA-approved alkaloids, dihydroergotamine and ergotamine, and a non-FDA-approved putative MAPKAPK2 inhibitor, ZINC73240059, as small molecule binders to the M1 region

Virtual screening by molecular docking was performed using 2,115 FDA-approved and 515,454 non-FDA-approved small molecules from the ZINC15 database ^38^ against residues of the internal mitochondrial sequence, M1 (aa: 35–41) in the N-terminal domain (NTD) of TDP-43. The X-ray crystallographic structure of the N-terminal domain, 5MDI (aa: 2–79), was used as the macromolecular structure used, with the grid box centered on the M1 residues. The top-ranking hits from virtual screening of 2,115 FDA-approved small molecules against the M1 region of the N-terminal domain of TDP-43 exhibited docking binding free energies ranging from −8.2 to −7.5 kcal/mol (**Supplementary Figure SF6a**). Among these, ZINC1612996 (Irinotecan) achieved the best docking score of −8.2 kcal/mol, followed by ZINC169289767 (Trypan Blue), three tautomeric forms of ZINC3978005 (Dihydroergotamine), ZINC164760756 (Simeprevir), and ZINC52955754 (Ergotamine). Dihydroergotamine and ergotamine are alkaloids used to treat migraines. ^41^ Irinotecan is an anti-cancer medication used to treat colon cancer and small-cell lung carcinoma by acting as a topoisomerase I inhibitor. ^42^ Simeprevir, a hepatitis C virus infection medication, is a serine protease inhibitor. ^43^

Among the 515,545 non-FDA-approved molecules in the category annotated and not for sale subset of the ZINC15 database, 98 had docking scores less than −8.5 when screened against the M1 region of the NTD of the TDP43 structure (**Supplementary Figure SF6b**). ZINC73240063 ranked top with a docking score of −9.4 kcal/mol. The next top molecules in this subset are ZINC45201764, ZINC73240059, ZINC45210931, ZINC2942260, and ZINC28707596, in the order of docking score. ZINC73240063 and ZINC73240059 in the above list are reported to be inhibitors of mitogen-activated protein kinase-activated protein kinase 2 (MAPKAPK2). ZINC45201764 is predicted to target mitogen-activated protein kinase 13. ^44^

The top-ranked molecules from the virtual screening of the FDA-approved and non-FDA-approved small-molecule subsets were subsequently blindly docked to NTD monomer taken out of the crystal structure of the dimeric NTD of TDP-43 (PDB ID: 6T4B). The docking using AutoDock 4.2 reported only three compounds, ZINC3978005 (Dihydroergotamine), ZINC52955754 (Ergotamine), and ZINC73240059, to interact with the M1 region (**Table 2**). These three molecules were subsequently further docked with the modeled full-length TDP-43 structure to determine their domain- and region-specificity. The RMSD values for the full-length and solved structures of various TDP-43 domains are provided in the **Supplementary Table S2**. Three of these molecules, ZINC3978005, ZINC52955754, and ZINC73240059, interacted at the binding pocket containing the M1 residues, while ZINC45210931, a top molecule from the virtual screening, interacted closely around M1 but not with the M1 residues (**Figure 4**).

**Figure 4:**
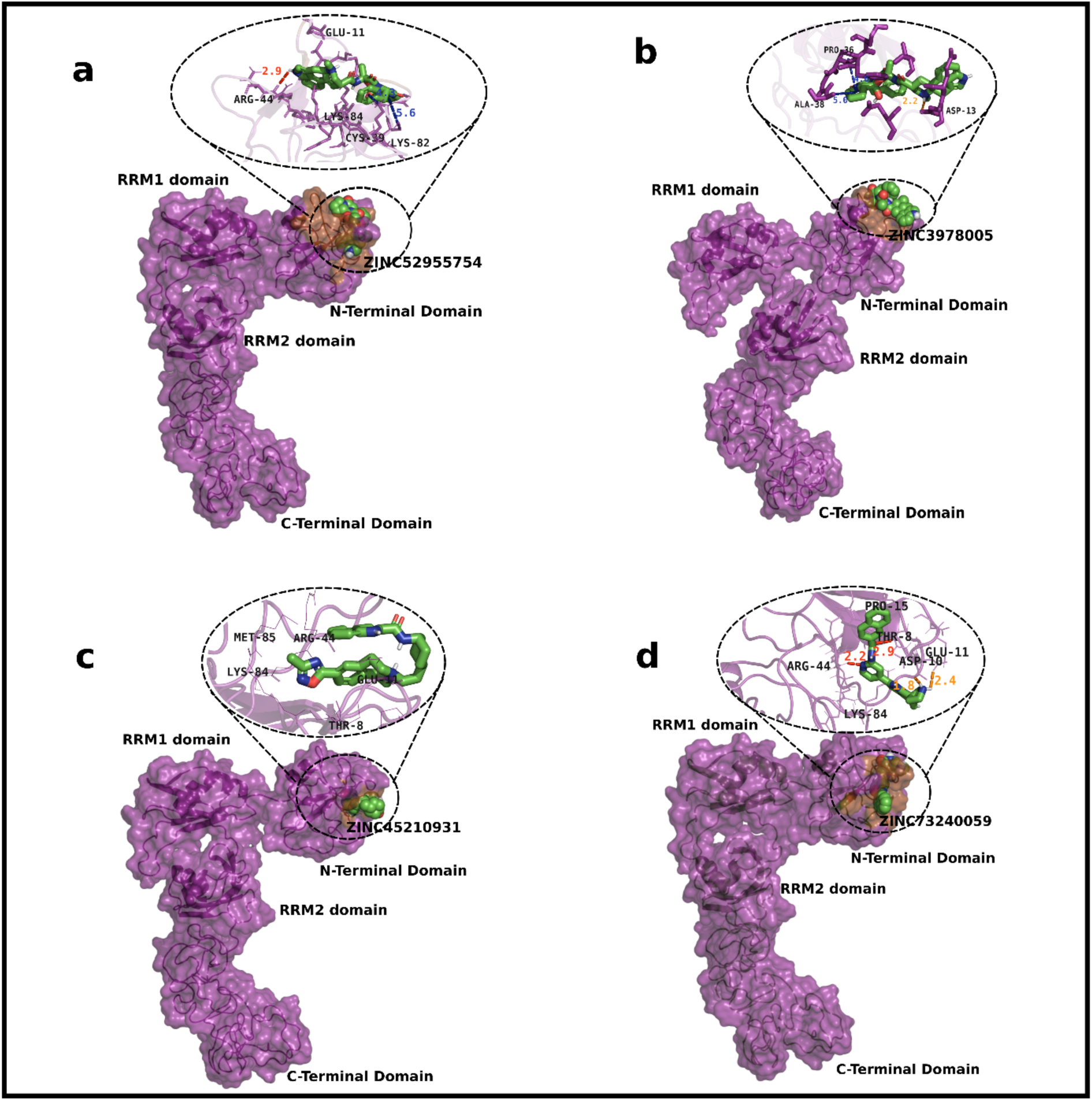
Docked poses of the small molecules with the I-TASSER-modeled structure of full-length TDP-43. **a)** The docked pose of the small molecule ZINC52955754 (Ergotamine) with full-length TDP-43 structure modeled using the online webserver ITASSER-MTD. **b)** The top-ranked docked pose of the small molecule ZINC3978005 (Dihydroergotamine) with the modeled full-length TDP-43 structure. **c)** The docked pose of the molecule ZINC45210931 with modeled full-length TDP-43 structure. **d)** The complex structure after blind docking for the molecule ZINC73240059 with the full-length modeled TDP-43 structure. The protein structure in the docked poses above is represented as New Cartoon and Surface representations in purple, with the M1 region (aa: 35–41) colored brown. The small molecules are represented as van der Waals’ spheres. A close view of the docking interactions is shown in the circle adjacent to the complete view of the docked structures. The blue color bonds represent pi-alkyl interactions. The hydrogen and cation-pi interactions are colored red and yellow, respectively.

**Table 2:**
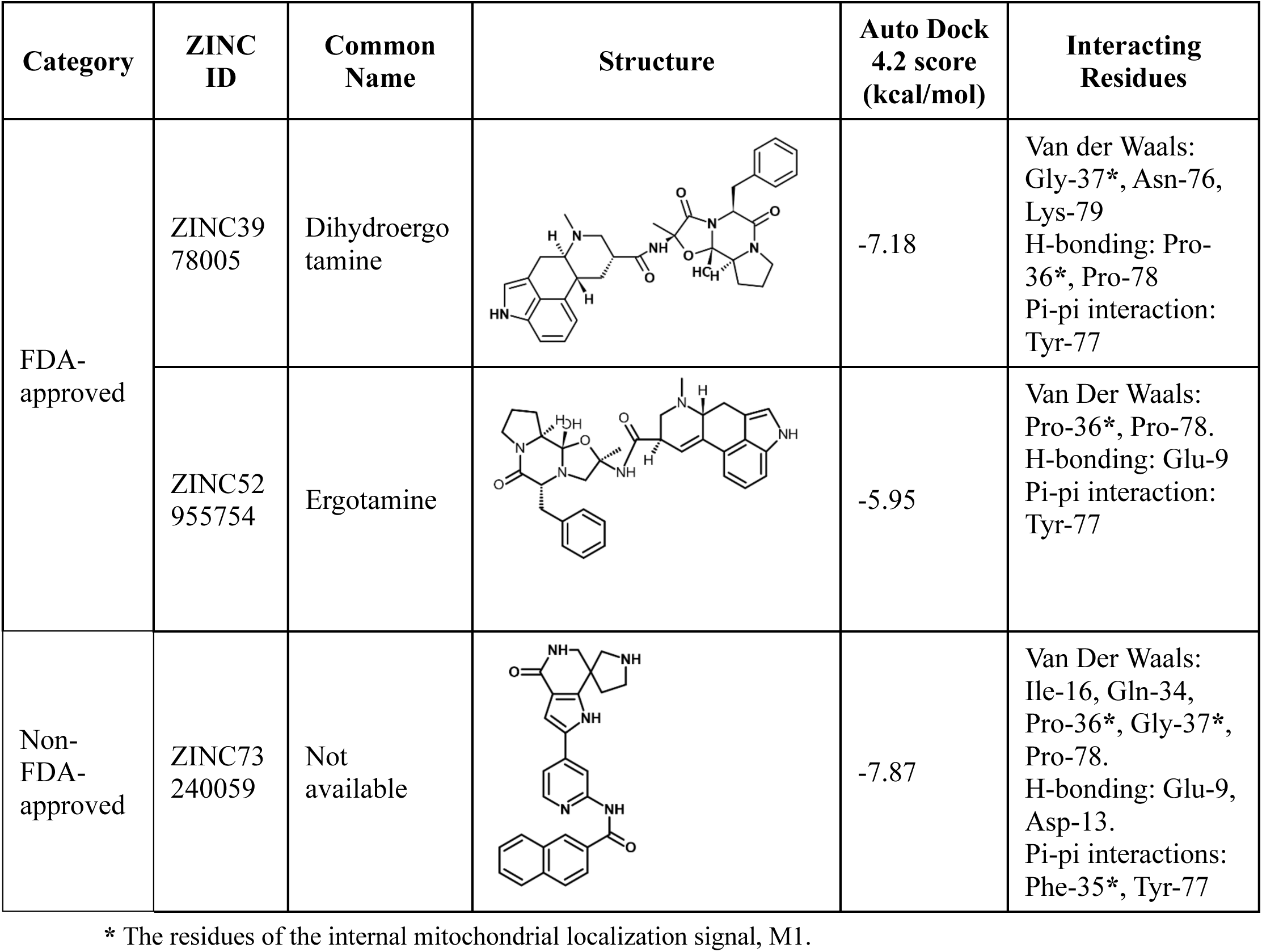
Top-ranked molecules from virtual screening against the N-terminal domain structure (PDB ID: 2N4P) of TDP-43.

| Category | ZINC ID | Common Name | Structure | Auto Dock 4.2 score (kcal/mol) | Interacting Residues |
| --- | --- | --- | --- | --- | --- |
| FDA-approved | ZINC3978005 | Dihydroergotamine |  | -7.18 | Van der Waals: Gly-37*, Asn-76, Lys-79<br>H-bonding: Pro-36*, Pro-78<br>Pi-pi interaction: Tyr-77 |
|  | ZINC52955754 | Ergotamine |  | -5.95 | Van Der Waals: Pro-36*, Pro-78.<br>H-bonding: Glu-9<br>Pi-pi interaction: Tyr-77 |
| Non-FDA-approved | ZINC73240059 | Not available |  | -7.87 | Van Der Waals: Ile-16, Gln-34, Pro-36*, Gly-37*, Pro-78.<br>H-bonding: Glu-9, Asp-13.<br>Pi-pi interactions: Phe-35*, Tyr-77 |

\* The residues of the internal mitochondrial localization signal, M1.

For a small molecule to become a *bona fide* binder to M1 for inhibition of the TDP-43’s mitochondrial localization, it should also be able to stably interact with M1 in the partially unfolded conformations of NTD which would expectedly prevent the interaction and recognition of M1 by the mitochondrial transporters. Thus, the intermediate conformations, obtained by clustering the trajectories at 50-250 ns intervals from the all-atom MD simulation of unfolding of TDP43-NTD (PDB ID: 2N4P), as described in the previous section, were also used to determine the interaction efficiency of the top-most screened candidate molecules on these intermediate conformations. Strikingly, the ZINC73240059 molecule (a putative MAPKAPK2 inhibitor) remained bound at the M1 region in greater number of these conformations in comparison to the ZINC3978005 molecule (Dihydroergotamine) (**Figure 5a-f**). Although ZINC73240059 and ZINC3978005 interacted with M1 in different conformations, the ZINC73240059 molecule preferred M1 for binding across most conformations in the unfolding trajectories that too with comparable docking scores (**Supplementary Table S3**).

**Figure 5:**
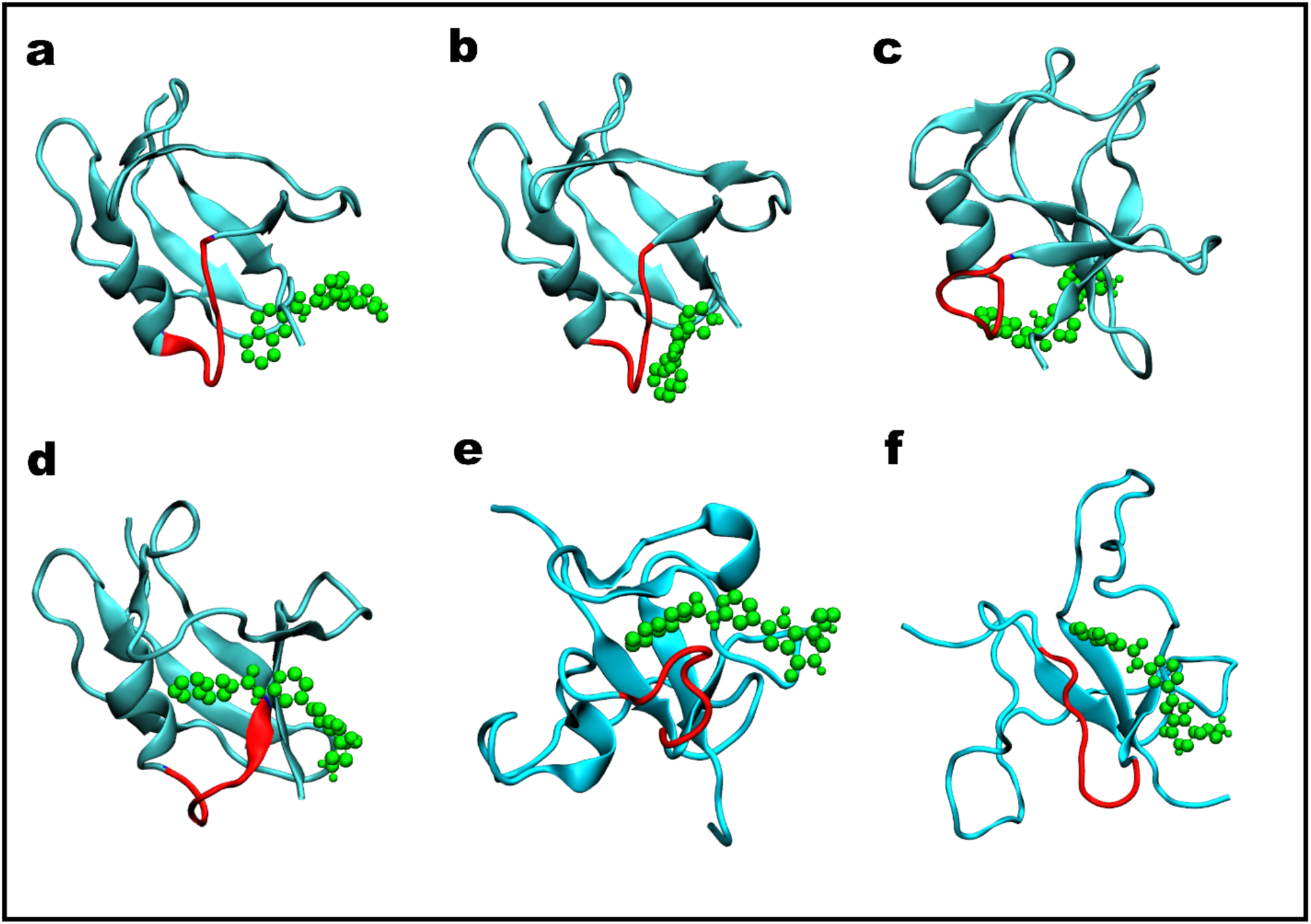
Docked poses of the ZINC73240059 with the intermediate structures obtained from unfolding of the N-terminal domain structure (PDB ID: 2N4P) of TDP-43. **a-e)** Representative average structures obtained from clustering of the 2N4P unfolding MD simulation with a 50 ns window interval until 300 ns, and whichever docked with ZINC73240059 (0-50, 50-100, 150-200, 200-250, 250-300 ns, respectively). **f)** A representative structure obtained from clustering of the 2N4P unfolding simulation trajectory from 300-500 ns docked with ZINC73240059. The protein macromolecule is shown in a new cartoon representation in cyan, with the M1 region (aa 35-41) marked in red. The small molecule ZINC73240059 is represented in green spheres.

Taken together, the ZINC73240059 molecule docked consistently to the M1 region of the two NTD structures of TDP-43 as well as the modeled full-length TDP-43 structure. The molecule interacted with the aromatic side chain of Phe-35 *via* a pi-pi interaction in addition to hydrophobic interactions with hydrophobic residues in the M1 region, making this region a preferred binding site. As the structural snapshots from the unfolding MD simulations indicate that ZINC73240059 can interact well with M1 in both folded as well as partially unfolded conformations of NTD therefore it is a suitable candidate molecule for further investigation. Hence, the ZINC73240059 molecule, and for comparison, other screened molecules like ZINC3978005, were furthermore investigated using MD simulations to evaluate their binding stabilities to the M1 residues of NTD.

### MD simulations corroborate stable binding of the ZINC73240059 molecule exclusively to the mitochondrial localization sequence, M1, of TDP-43

All-atom MD simulations were performed on the screened molecules to assess the stability of their binding during protein dynamics. Top-ranked molecules bound to the NTD of TDP-43 from the previous section were considered as the starting structure, and the all-atom MD simulations were performed in the explicit solution as mentioned in **Table 3**. Two molecules, namely the FDA-approved Dihydroergotamine (ZINC3978005) and the non-FDA-approved ZINC73240059, showed stable interactions with the N-terminal domain of TDP-43 as compared to the other ligands that topped the virtual screening including ZINC13556029, ZINC45210931 considered for evaluation from the small-molecule library (**Supplementary Figures SF7 and SF8**). Two molecules from the non-FDA-approved subsets used in the virtual screening were further used as control molecules for the MD simulation strategy. ZINC45210931, a molecule that topped in the virtual screening, but not consistently in other docking strategies, was evaluated against the 5MDI and 2N4P structures of NTD. Another control molecule, ZINC13556029, that showed a docking score of −4.5 Kcal/mol in the virtual screening was also investigated against 2N4P to check if the molecule could interact with M1 in the flexible protein conformations during MD simulations. Both of these molecules did not consistently interact with the N-terminal domain structures therefore further MD simulation analysis were performed only with the ZINC73240059 molecule from the library of non-FDA-approved molecules (**Supplementary Figures SF7** and **SF8**).

**Table 3:** Details of the MD simulation system containing the complex of the screened molecules and the N-terminal domain (NTD) of TDP-43.

| NTD Protein Structure (PDB ID) | Ligand | Number of atoms in the system* |
| --- | --- | --- |
| 5MDI | None (Apo) | 13185<sup>#</sup> |
|  | ZINC73240059 | 13238<sup>#</sup> |
|  | ZINC45210931 | 13227<sup>#</sup> |
| 2N4P | None (Apo) | 21,571 |
|  | ZINC3978005 (Dihydroergotamine) | 21,576 |
|  | ZINC73240059 | 21,387 |
|  | ZINC13556029 | 21,596<sup>#</sup> |
|  | ZINC45210931 | 21,637<sup>#</sup> |
| 6T4B | None (Apo) | 21,403<sup>#</sup> |
|  | ZINC3978005 (Dihydroergotamine) | 22,310<sup>#</sup> |
|  | ZINC73240059 | 21,477<sup>#</sup> |
|  | ZINC52955754 (Ergotamine) | 21,472<sup>#</sup> |
|  | ZINC64033452 | 21,438<sup>#</sup> |

\* Number of atoms in the system = Atoms of Protein + Ligand + Solvent Water + Ion (Sodium + Chlorine) added to neutralize and make the final salt concentration of the solvent to 0.15 M. <sup>#</sup> Results corresponding to these simulations can be found in supplementary information.

Visualizing the 500 ns MD simulation of these top molecules docked to the protein showed that the molecule ZINC73240059 remains bound to the M1 region of the NTD NMR structure (PDB ID: 2N4P, aa: 1-77), while Dihydroergotamine (ZINC3978005) did not remain stably bound (**<u>Supplementary Movie S2</u>**). RMSD of the solution NMR structure of the N-terminal of TDP-43 (PDB ID: 2N4P) in the presence of ZINC73240059 converged at ∼2.5 Å, while the RMSD in the apo structure converged at ∼3.5 Å. Similarly, RMSD of the protein in 2N4P + ZINC3978005 complex also converged at ∼2.5 Å (**Supplementary Figure SF9a**). Characterized using RMSF analysis, the protein in complex with ZINC73240059 exhibited insignificant conformational fluctuations in the amino acid residues near the M1 region, where it binds, as compared to the protein alone RMSF (**Supplementary Figure SF9b**). The average interaction profile from simulations (hydrogen bonds and other non-covalent) showed that residues of M1 interact with ZINC73240059 for most of the simulation time, along with other residues of the N-terminal domain of TDP-43, unlike Dihydroergotamine (ZINC3978005) (**Figures 6a and b**). The binding free energy of these two small molecules, computed using the Molecular Mechanics/Poisson-Boltzmann Surface Area (MMPBSA) method for the last 300 ns of MD simulation (**Figures 6c and d**), indicates their affinity to the protein during dynamics. The MMPBSA analysis suggests that ZINC73240059 interacts relatively strongly at M1 of the N-terminal domain, with an average binding free energy of −18.76 ± 0.25 kcal/mol, the lowest among the ligands investigated. (**Supplementary Table S4**)

**Figure 6:**
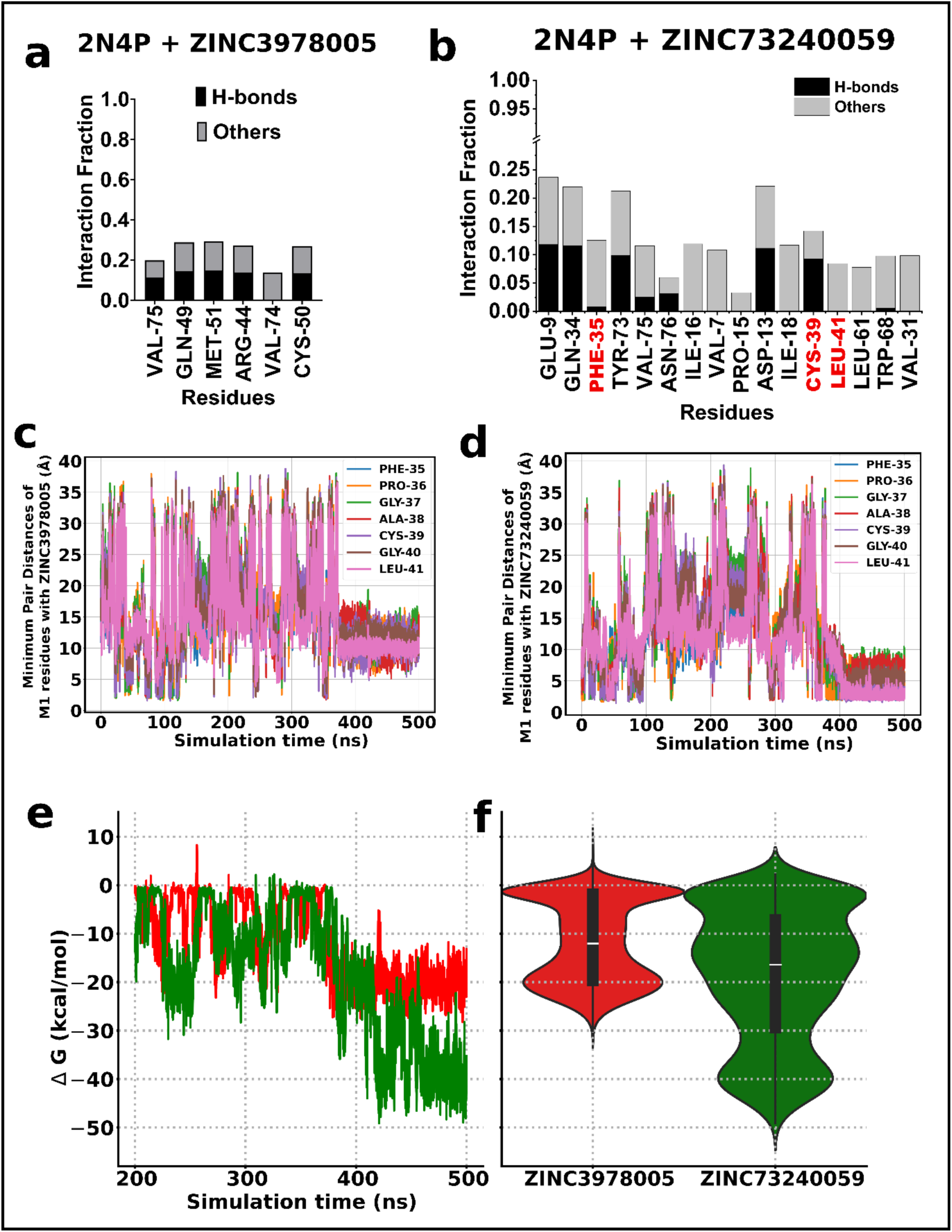
MD simulations of the virtually screened compounds with the solution NMR structure (2N4P) of the N-terminal domain of TDP-43. **a-b)** The amino acid residue-wise interaction fraction profile of the TDP-43 NTD structure, 2N4P, with the small molecules Dihydroergotamine (ZINC3978005) and ZINC7340059 obtained from the analysis of 500 ns MD simulations. The residues of the mitochondrial localization sequence, M1 (aa: 35-41), are shown in red. An interaction fraction value of “1” means that the residue of the protein interacts with the small molecule with both the hydrogen bond and other non-covalent interactions throughout the simulation time (H-bond + Other bonds = 0.5 + 0.5 = 1.0). Minimum distances between the small molecule and the M1 amino acid residues with **c)** ZINC3978005 and **d)** ZINC73240059 over the simulation time of 500 ns. **e-f)** MMPBSA binding free energy profile for the ZINC3978005 and ZINC73240059 with TDP-43 NTD structure, 2N4P, shown as a line plot in the left panel and the probability distribution of the values as a violin plot in the right panel.

In another set of MD simulations with the X-ray crystallographic structure (PDB ID: 6T4B, aa: 2-79) of the N-terminal domain of the TDP-43, the protein complex shows lower RMSD with ZINC73240059, ZINC3978005 (Dihydroergotamine), ZINC52955754 (Ergotamine) and a control molecule which showed lower affinity in docking (ZINC64033452), similar to the simulations with the complex with NMR NTD structure (**Supplementary Figure SF10a**). The RMSF of the protein and drug complex simulations showed fluctuations in residues 20-25 and 40-48 for dihydroergotamine, as compared to the protein alone RMSF for 6T4B (**Supplementary Figure SF10b**).

Dihydroergotamine (ZINC3978005) interacted with the residues that are not from M1 for most of the simulation time (**Supplementary Figure SF11a**), whereas ZINC73240059 interacted with the M1 region (**Supplementary Figure SF11b**). These two molecules, ZINC73240059 and ZINC3978005, gave an average free energy value of −14.24 kcal/mol and −14.51 kcal/mol with 6T4B, respectively, for the last 300 ns simulation trajectory as shown in **Supplementary Figure SF11c, d** and **Supplementary Table S4**. Furthermore, the per-residue decomposition of the free energy change for the 6T4B+ZINC73240059 complex (Supplementary **Table S5),** identified contribution from the M1 residues, unlike for 6T4B+ZINC3978005 complex, confirming the stable interaction with M1.

As corroborated with the observations from the virtual screening, blind docking and MD simulations, the putative MAPKAPK2 inhibitor molecule ZINC73240059 (N-[4-(1′,4′,5′,6′-Tetrahydro-4′-oxospiro[pyrrolidine-3,7′-[7H]pyrrolo[3,2-c]pyridin]-2′-yl)-2-pyridinyl]-2-naphthalenecarboxamide) interacts strongly and preferentially with the internal mitochondrial sequence, M1, of TDP-43. Thus, we propose this molecule as a potential inhibitor of the M1-mediated aberrant mitochondrial localization of TDP-43 which may be investigated in future in relevant disease models.

## 3.0 Conclusion

Previous studies found certain sequences within the TDP-43 protein, termed M1, M3, and M5, that can facilitate aberrant mitochondrial localization of TDP-43, and inhibition of the TDP-43’s mitochondrial import either by deleting these amino acid stretches or by competitive inhibition could alleviate the TDP-43-related toxicity. The TDP-43 import inhibition, thus far, has been obtained only by certain peptides that mimicked these mitochondrial localization sequences and hence functioned as competitive inhibitors thereby preventing the TDP-43’s import ^13^. Hence, directly targeting these mitochondrial localization sequences by binding of small molecules with high affinity, appears as an attractive strategy to prevent the mitochondrial localization of TDP-43 and its consequent toxicity. However, some of these sequences such as M1, are largely solvent inaccessible, therefore, how they interact with the mitochondrial import machinery to facilitate TDP-43’s transit to mitochondria is unclear. Thus, elucidating their conformational dynamics can provide useful insights into their mechanism and facilitate targeting them with small molecules. In this study, we illustrate *in silico* unfolding of the N-terminal domain (NTD) of TDP-43 and reveal the disruption of hydrogen-bonded interactions involving an M1 residue at the end of α1 (Phe-35) and among residues of the β4-β5 bridge as initial events in the unfolding of NTD in addition to the highly fluctuating β3 residues reported earlier. ^23,24^ We visualize the unfolding of NTD of TDP-43 as a potential mimic for the exposure of the M1 residues to the mitochondrial import machinery, using MD simulations performed on two separate PDB structures of NTD, the solution NMR structure, 2N4P, and the X-ray crystallographic structure, 6T4B. As small molecules with better pharmacokinetic properties compared to the peptides ^45^ could also be potential inhibitors to the mitochondrial import of TDP-43 therefore, we performed a large-scale virtual screening of small molecules to find potential molecules that can interact with the mitochondrial signal M1 (aa 35–41) of TDP-43 that may possibly in future be investigated to inhibit TDP-43’s mitochondrial import. This large-scale virtual screening using docking techniques and MD simulations in explicit solvent at 300K and 310K identified one molecule, ZINC73240059, that interacts stably with the M1 region. This molecule showed consistent and stable binding to the M1 region of both the solution NMR structure (2N4P) and the X-ray crystallographic structures (5MDI and 6T4B) of NTD of TDP-43. The average free energies for the binding of ZINC73240059 to 6T4B and 2N4P structures of the NTD for the last 300 ns of the 500 ns MD simulations was respectively calculated to be –14.24 ± 0.1 kcal/mol and −18.76 ± 0.25 kcal/mol. The per-residue contribution to the change in the free energy of the complex of ZINC73240059 with 6T4B showed that Pro-36 of M1 and residues Tyr-77 and Pro-78 in the region around M1 showed stable interactions. Overall, based on the analysis of equilibrium MD simulation data, we propose a mechanism of unfolding of NTD in which the interactions involving the M1 residue (Phe-35) and those among the β4-β5 strands are initially disrupted driving the unfolding of NTD and a putative MAPKAPK2 inhibitor molecule, ZINC73240059, as a stable binder to M1 with a potential to inhibit the aberrant mitochondrial mislocalization of TDP-43.

## Methodology

### System preparation

All-atom MD simulation in the current study was performed on three N-terminal domain structures. Among the three N-terminal domain structures were a solution NMR structure (PDB ID: 2N4P, aa: 1-77) with a 6x histidine tag. ^35^ The histidine tag and additional sequences not belonging to TDP-43 were removed from the 2N4P prior to docking and simulation experiments. Other N-terminal domain structures used in the study include X-ray crystallographic structures (PDB ID: 6T4B, aa: 2-79) ^37^ and (PDB ID: 5MDI, aa: 2-79). ^36^ In unfolding simulations, urea was used as a denaturant. The urea structure was downloaded from the CHARMM small-molecule library on the CHARMM-GUI web server. ^46^ The GROMACS-compatible topology for urea was later created using the CHARMM General Force Field (CGenFF v4.0). ^47–49^ Similarly, for the protein-ligand simulations, the topology for the ligand from the docked complex was also generated using the CGenFF v4.0 web server.

### Simulation protocol

All-atom MD simulations were performed on the Groningen Machine for Chemical Simulation (GROMACS) v. 21.4. GROMACS is a GNU-licensed software for MD simulations. ^50^ The protein topology was prepared using the built-in pdb2gmx command with the CHARMM36m-force field. ^51^ The TIP3P-modified water solvent model ^52^, placed in a dodecahedron box with edges 15 Å away from the protein, was used to solvate the protein. An in-built *gmx insert-molecule* command was used to insert the urea molecule at the desired concentration into simulations that require it. The solvated systems were supplemented with Na^+^ and Cl^−^ counterions to achieve a 150 mM NaCl concentration, after neutralizing the system using a genion script in GROMACS. Thus, the assembled system containing the protein, water, ions, and, where applicable, the denaturant was subjected to energy minimization using the steepest descent algorithm to avoid atom clashes in the built configuration. NVT and NPT equilibration at the temperatures corresponding to the different simulation conditions, each for 10ns, was performed after energy minimization with harmonic position restraints applied to the protein’s heavy atoms. An unrestrained production run for the equilibrated system was performed with a 2-fs integration time step at different temperature conditions. Periodic boundary conditions were applied along the x-, y-, and z-axes. Long-range electrostatic interactions were calculated using the Particle Mesh Ewald (PME) method with a resolution grid of 1.2 Å. ^53^ The temperatures and 1 bar pressure for the equilibration simulations were maintained using the V-rescale and Berendsen coupling methods ^54^, while 1 bar isotropic pressure in the unrestrained production run was maintained using the Parrinello-Rahman barostat. ^55^ For protein-ligand complex simulations, the combined topology of the protein-ligand complex was provided as input to GROMACS. The initial snapshot was obtained through the docking of the molecules to protein.

### Simulation Analysis

In-built gmx scripts for the (root-mean-squared deviations) RMSD, root-mean-squared fluctuations (RMSF), and pair-distance calculations were used. Residue-wise solvent accessibility was calculated using a Tcl/Tk script in Visual Molecular Dynamics (VMD). ^56^ MDTraj is an open-source Python library for analyzing and manipulating the MD trajectories. ^57^ It was employed to determine the fraction of native contacts (Q). The fraction of native contacts measures the relative number of atom pairs present at a particular trajectory frame compared to the total number of atom pairs present in the reference (native) structure, as defined earlier. ^58^ The change in free energy (ΔG) as a function of conformational state parameters was calculated using Boltzmann inversion as follows:

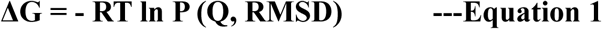

R = Universal gas constant (8.314 J mol^−1^ K^−1^)

T = Temperature (K)

P (Q, RMSD) = Joint probability of fraction of native contacts (Q) and RMSD.

The free energy change is normalized to the minimum value obtained; hence, the lowest energy corresponds to zero (0), as calculated in earlier works. ^14,15^ The minimum pair distances were calculated using the *gmx pairdist* command among the residue pairs previously reported to form hydrogen bonds. ^35^ The centre of mass of the residues is used as the reference and output selection positions for the distance calculation. The folded native state was measured to study the sequence of events leading to the unfolding of the N-terminal structure at high temperatures and in the presence of urea. For the protein-ligand complex simulations, the lifetime of a residue-ligand interaction in the trajectory was computed using an in-house Python script. The input to the script consisted of the interaction type and the interaction scores for each residue, as obtained from PyContact v1.0.5, a GUI-based tool for evaluating non-covalent interactions. ^59^ The compressed trajectory and topology files from the protein-ligand simulations were provided as input to PyContact, along with a stride value of 100. The default criteria of 2.5Å and 5Å for H-bonds and other non-covalent interactions were considered. The 120° angle cut-off was also applied to H-bond interactions. The molecular mechanics Poisson-Boltzmann surface area (MM-PBSA) continuum method was employed to calculate the free energy change for the complex and its per-residue contributions using the gmx_MMPBSA tool v.1.6.2. ^60^ A single-trajectory MMPBSA analysis was performed with the complex trajectory given as input. 5,001 frames per trajectory were submitted, with a stride of 100 ps over the entire 500 ns simulation used in the study, which contained 50,000 frames. The internal and external dielectric constants of 1.0 and 80.0, respectively, were used, with the fill ratio and spacing for the PB calculation set to 4.0 and 2 Å, respectively. The implicit solvent model, with an ionic strength of 0.15 M and a distance cutoff of 6 Å, was used to calculate the free energy and per-residue contributions. GraphPad Prism 10.3.0.507 was used for the interaction fraction and change in free energy plots. The 3D representations of the macromolecule and ligand were created using VMD and PyMol. ^61^

### Modeling of the full-length structure

The full-length structure of TDP-43 has not been resolved experimentally yet; hence, the modeling of the full-length structure was performed on I-TASSER-MTD. I-TASSER-MTD is a free web server for predicting multidomain protein structures and function prediction. ^62^ The complete sequence of the TDP-43 in the FASTA format from Uniprot (ID: Q13148_1) was provided as input to the web server for prediction. The first model from the web server was used for molecular docking.

### System preparation for virtual screening

The small-molecule subsets used in the structure-based virtual screening of the current study comprise FDA-approved compounds (2,115) and non-approved compounds currently under research for various diseases (515,545) from the ZINC15 library (https://zinc15.docking.org). ZINC15 is a publicly available database of small molecules that allows easy retrieval of 3D structures via its tranche browser. ^38^ The small molecules were downloaded in MOL2 format. They were converted to the PDBQT format using Open Babel v3.1.0, a GNU-licensed chemical toolbox, making them compatible with the docking software used in the study. ^63^ The preparation steps required to dock the ligand, such as computing Gasteiger partial charges for each atom and adding explicit hydrogen atoms, were performed in Open Babel during the conversion. The target of the virtual screening was the internal mitochondrial signal M1 in the N-terminal domain of TDP-43, as observed in the crystal structures (PDB ID: 5MDI, aa: 2-79). A publicly available Perl script performed large-scale virtual screening of small molecules by defining a grid box around the M1 residue (aa: 35-41) as the search space, which was provided as an input parameter for AutoDock Vina v1.12 docking. AutoDock Vina is an open-source molecular docking program. ^64^ Other parameters were supplied to the script via a configuration file, which included the protein molecule (5MDI) as the receptor. The Perl script automatically considers the ligand files as sequential inputs.

### Blind molecular docking protocol

The top molecules from the virtual screening were docked against the crystal structure of the N-terminal domain of TDP-43 (PDB ID: 6T4B), the solution NMR structure (PDB ID: 2N4P), the modeled full-length structure, or the snapshots from the unfolding simulation trajectories. The small molecules and macromolecular structures for each docking were prepared as described in the previous section, except that the grid box was extended to cover the entire macromolecule. The blind docking for the top-scoring molecules in all the above-mentioned cases was performed using AutoDock 4.2.6. AutoDock 4.2.6 is GNU-licensed docking software that requires an autogrid program to precalculate grid maps for the atoms involved using the MGL tools v1.5.7 software suite. ^65^ A Lamarckian genetic algorithm (GA) was used for 50 runs as a search parameter in docking performed on the modeled full-length TDP-43.

## Acknowledgements

We thank IIT Hyderabad, funded by the Ministry of Education (MoE), Govt. of India, for the research infrastructure and support. RB thanks MoE, Govt. of India, for Prime Minister Research Fellowship (PMRF) (PMRF ID: 2001298). SB, and JB thanks MoE, Govt. of India, for JRF. H.J. would like to acknowledge the startup research grant (SRG/ 2022/002109) by Science and Engineering Research Board (India) and Inspire faculty fellowship (IFA-20-PH-256) by Department of Science of Technology (Government of India) for supporting this work. Supercomputer time was generously provided by NSM through the PARAM Seva machine at IIT Hyderabad. Basant K Patel thanks SERB-DST, Govt. of India for a research grant (Grant no: SERB/CRG/2021/006856)

## CRediT author contribution statement

**RB**: Writing – review & editing, Writing – original draft, Investigation.

**SB**: Writing – review & editing, Writing – original draft, Investigation.

**JB**: Writing – review & editing, Investigation.

**HJ**: Writing – review & editing, Supervision, Project administrations, Investigations, Conceptualization.

**BKP**: Writing – reviewing & editing, Supervision, Project administration, Conceptualization.

## References

(1) Prasad, A.; Bharathi, V.; Sivalingam, V.; Girdhar, A.; Patel, B. K. Molecular Mechanisms of TDP-43 Misfolding and Pathology in Amyotrophic Lateral Sclerosis. Front. Mol. Neurosci. 2019, 12, 25. 10.3389/fnmol.2019.00025.

(2) Roy, A.; Chhetry, S.; Deka, H.; Roy, R.; Bhattacharya, P.; Patel, B. K.; Borah, A. TDP-43: A Critical Amplifier of Alzheimer’s Disease beyond Amyloid and Tau. Neuroscience 2026, 608, 178–189. 10.1016/j.neuroscience.2026.05.024.

(3) François-Moutal, L.; Perez-Miller, S.; Scott, D. D.; Miranda, V. G.; Mollasalehi, N.; Khanna, M. Structural Insights Into TDP-43 and Effects of Post-Translational Modifications. Front. Mol. Neurosci. 2019, 12, 301. 10.3389/fnmol.2019.00301.

(4) Sreedharan, J.; Blair, I. P.; Tripathi, V. B.; Hu, X.; Vance, C.; Rogelj, B.; Ackerley, S.; Durnall, J. C.; Williams, K. L.; Buratti, E.; Baralle, F.; De Belleroche, J.; Mitchell, J. D.; Leigh, P. N.; Al-Chalabi, A.; Miller, C. C.; Nicholson, G.; Shaw, C. E. TDP-43 Mutations in Familial and Sporadic Amyotrophic Lateral Sclerosis. Science 2008, 319 (5870), 1668–1672. 10.1126/science.1154584.

(5) Neumann, M.; Sampathu, D. M.; Kwong, L. K.; Truax, A. C.; Micsenyi, M. C.; Chou, T. T.; Bruce, J.; Schuck, T.; Grossman, M.; Clark, C. M.; McCluskey, L. F.; Miller, B. L.; Masliah, E.; Mackenzie, I. R.; Feldman, H.; Feiden, W.; Kretzschmar, H. A.; Trojanowski, J. Q.; Lee, V. M.-Y. Ubiquitinated TDP-43 in Frontotemporal Lobar Degeneration and Amyotrophic Lateral Sclerosis. Science 2006, 314 (5796), 130–133. 10.1126/science.1134108.

(6) Janssens, J.; Kleinberger, G.; Wils, H.; Van Broeckhoven, C. The Role of Mutant TAR DNA-Binding Protein 43 in Amyotrophic Lateral Sclerosis and Frontotemporal Lobar Degeneration. Biochem. Soc. Trans. 2011, 39 (4), 954–959. 10.1042/BST0390954.

(7) Buratti, E. Targeting TDP-43 Proteinopathy with Drugs and Drug-like Small Molecules. Br. J. Pharmacol. 2021, 178 (6), 1298–1315. 10.1111/bph.15148.

(8) Prasad, A.; Raju, G.; Sivalingam, V.; Girdhar, A.; Verma, M.; Vats, A.; Taneja, V.; Prabusankar, G.; Patel, B. K. An Acridine Derivative, [4,5-Bis{(N-Carboxy Methyl Imidazolium)Methyl}acridine] Dibromide, Shows Anti-TDP-43 Aggregation Effect in ALS Disease Models. Sci. Rep. 2016, 6 (1), 39490. 10.1038/srep39490.

(9) Girdhar, A.; Bharathi, V.; Tiwari, V. R.; Abhishek, S.; Deeksha, W.; Mahawar, U. S.; Raju, G.; Singh, S. K.; Prabusankar, G.; Rajakumara, E.; Patel, B. K. Computational Insights into Mechanism of AIM4-Mediated Inhibition of Aggregation of TDP-43 Protein Implicated in ALS and Evidence for in Vitro Inhibition of Liquid-Liquid Phase Separation (LLPS) of TDP-432C-A315T by AIM4. Int. J. Biol. Macromol. 2020, 147, 117–130. 10.1016/j.ijbiomac.2020.01.032.

(10) Meshram, V. D.; Balaji, R.; Saravanan, P.; Subbamanda, Y.; Deeksha, W.; Bajpai, A.; Joshi, H.; Bhargava, A.; Patel, B. K. Computational Insights Into the Mechanism of EGCG’s Binding and Inhibition of the TDP-43 Aggregation. Chem. Biol. Drug Des. 2024, 104 (4), e14640. 10.1111/cbdd.14640.

(11) Preethi, S.; Bharathi, V.; Patel, B. K. Zn2+ Modulates in Vitro Phase Separation of TDP-432C and Mutant TDP-432C-A315T C-Terminal Fragments of TDP-43 Protein Implicated in ALS and FTLD-TDP Diseases. Int. J. Biol. Macromol. 2021, 176, 186–200. 10.1016/j.ijbiomac.2021.02.054.

(12) Saravanan, P.; Bharathi, V.; Veerabhadraswamy, P.; Patel, B. K. Enhanced *in Vitro* Aggregation, but Not Phase Separation, of TDP-43 and Its C-Terminal Fragments Generates Intrinsic Deep-Blue Autofluorescence. New J. Chem. 2025, 49 (38), 16669–16690. 10.1039/D5NJ02842F.

(13) Wang, W.; Wang, L.; Lu, J.; Siedlak, S. L.; Fujioka, H.; Liang, J.; Jiang, S.; Ma, X.; Jiang, Z.; Da Rocha, E. L.; Sheng, M.; Choi, H.; Lerou, P. H.; Li, H.; Wang, X. The Inhibition of TDP-43 Mitochondrial Localization Blocks Its Neuronal Toxicity. Nat. Med. 2016, 22 (8), 869–878. 10.1038/nm.4130.

(14) Lu, J.; Duan, W.; Guo, Y.; Jiang, H.; Li, Z.; Huang, J.; Hong, K.; Li, C. Mitochondrial Dysfunction in Human TDP-43 Transfected NSC34 Cell Lines and the Protective Effect of Dimethoxy Curcumin. Brain Res. Bull. 2012, 89 (5–6), 185–190. 10.1016/j.brainresbull.2012.09.005.

(15) Duan, W.; Li, X.; Shi, J.; Guo, Y.; Li, Z.; Li, C. Mutant TAR DNA-Binding Protein-43 Induces Oxidative Injury in Motor Neuron-like Cell. Neuroscience 2010, 169 (4), 1621–1629. 10.1016/j.neuroscience.2010.06.018.

(16) Bajpai, A.; Bharathi, V.; Patel, B. K. Therapeutic Targeting of the Oxidative Stress Generated by Pathological Molecular Pathways in the Neurodegenerative Diseases, ALS and Huntington’s. Eur. J. Pharmacol. 2025, 987, 177187. 10.1016/j.ejphar.2024.177187.

(17) Tanveer, S.; Bajpai, A.; Rajput, S. J.; Saravanan, P.; Patel, B. K. Targeting Oxidative Stress to Combat Neurodegenerative Diseases, Amyotrophic Lateral Sclerosis and Huntington’s, for Better Brain Health and Recovery. Curr. Opin. Physiol. 2025, 46, 100872. 10.1016/j.cophys.2025.100872.

(18) Bharathi, V.; Girdhar, A.; Prasad, A.; Verma, M.; Taneja, V.; Patel, B. K. Use of Ade1 and Ade2 Mutations for Development of a Versatile Red/White Colour Assay of Amyloid-Induced Oxidative Stress in Saccharomyces Cerevisiae. Yeast 2016, 33 (12), 607–620. 10.1002/yea.3209.

(19) Wang, P.; Deng, J.; Dong, J.; Liu, J.; Bigio, E. H.; Mesulam, M.; Wang, T.; Sun, L.; Wang, L.; Lee, A. Y.-L.; McGee, W. A.; Chen, X.; Fushimi, K.; Zhu, L.; Wu, J. Y. TDP-43 Induces Mitochondrial Damage and Activates the Mitochondrial Unfolded Protein Response. PLOS Genet. 2019, 15 (5), e1007947. 10.1371/journal.pgen.1007947.

(20) Ma, J.; Liu, L.; Song, L.; Liu, J.; Yang, L.; Chen, Q.; Wu, J. Y.; Zhu, L. Integration of FUNDC1-Associated Mitochondrial Protein Import and Mitochondrial Quality Control Contributes to TDP-43 Degradation. Cell Death Dis. 2023, 14 (11), 735. 10.1038/s41419-023-06261-6.

(21) Dudek, J.; Rehling, P.; Van Der Laan, M. Mitochondrial Protein Import: Common Principles and Physiological Networks. Biochim. Biophys. Acta BBA - Mol. Cell Res. 2013, 1833 (2), 274–285. 10.1016/j.bbamcr.2012.05.028.

(22) Yu, C.-H.; Davidson, S.; Harapas, C. R.; Hilton, J. B.; Mlodzianoski, M. J.; Laohamonthonkul, P.; Louis, C.; Low, R. R. J.; Moecking, J.; De Nardo, D.; Balka, K. R.; Calleja, D. J.; Moghaddas, F.; Ni, E.; McLean, C. A.; Samson, A. L.; Tyebji, S.; Tonkin, C. J.; Bye, C. R.; Turner, B. J.; Pepin, G.; Gantier, M. P.; Rogers, K. L.; McArthur, K.; Crouch, P. J.; Masters, S. L. TDP-43 Triggers Mitochondrial DNA Release via mPTP to Activate cGAS/STING in ALS. Cell 2020, 183 (3), 636–649.e18. 10.1016/j.cell.2020.09.020.

(23) Li, R.; Singh, R.; Kashav, T.; Yang, C.; Sharma, R. D.; Lynn, A. M.; Prasad, R.; Prakash, A.; Kumar, V. Computational Insights of Unfolding of N-Terminal Domain of TDP-43 Reveal the Conformational Heterogeneity in the Unfolding Pathway. Front. Mol. Neurosci. 2022, 15, 822863. 10.3389/fnmol.2022.822863.

(24) Prakash, A.; Kumar, V.; Meena, N. K.; Lynn, A. M. Elucidation of the Structural Stability and Dynamics of Heterogeneous Intermediate Ensembles in Unfolding Pathway of the N-Terminal Domain of TDP-43. RSC Adv. 2018, 8 (35), 19835–19845. 10.1039/C8RA03368D.

(25) Marzi, I.; Pieraccini, G.; Bemporad, F.; Chiti, F. Detection of an Intermediate in the Unfolding Process of the N-Terminal Domain of TDP-43. ACS Omega 2025, 10 (6), 5616–5633. 10.1021/acsomega.4c08617.

(26) Bharathi, V.; Girdhar, A.; Patel, B. K. Role of CNC1 Gene in TDP-43 Aggregation-Induced Oxidative Stress-Mediated Cell Death in S. Cerevisiae Model of ALS. Biochim. Biophys. Acta BBA - Mol. Cell Res. 2021, 1868 (6), 118993. 10.1016/j.bbamcr.2021.118993.

(27) Bajpai, A.; Bharathi, V.; Patel, B. K. Therapeutic Targeting of the Oxidative Stress Generated by Pathological Molecular Pathways in the Neurodegenerative Diseases, ALS and Huntington’s. Eur. J. Pharmacol. 2025, 987, 177187. 10.1016/j.ejphar.2024.177187.

(28) Balaji, R.; Joshi, H.; Patel, B. K. Elucidating the Conformational Dynamics of the Mitochondrial Localization Signal, M3, of TDP-43 and Accessing Potential Binders Using Molecular Docking and Simulation. Comput. Biol. Chem. 2026, 123, 109029. 10.1016/j.compbiolchem.2026.109029.

(29) Kitchen, D. B.; Decornez, H.; Furr, J. R.; Bajorath, J. Docking and Scoring in Virtual Screening for Drug Discovery: Methods and Applications. Nat. Rev. Drug Discov. 2004, 3 (11), 935–949. 10.1038/nrd1549.

(30) Hollingsworth, S. A.; Dror, R. O. Molecular Dynamics Simulation for All. Neuron 2018, 99 (6), 1129–1143. 10.1016/j.neuron.2018.08.011.

(31) Lindorff-Larsen, K.; Piana, S.; Dror, R. O.; Shaw, D. E. How Fast-Folding Proteins Fold. Science 2011, 334 (6055), 517–520. 10.1126/science.1208351.

(32) Fernando, K. S.; Chau, Y. Multi-Scale in Silico Analysis of the Phase Separation Behavior of FUS Mutants. J. Mater. Chem. B 2024, 12 (48), 12608–12617. 10.1039/D4TB01512F.

(33) Grasso, G.; Danani, A. Molecular Simulations of Amyloid Beta Assemblies. Adv. Phys. X 2020, 5 (1), 1770627. 10.1080/23746149.2020.1770627.

(34) Okumura, H. Perspective for Molecular Dynamics Simulation Studies of Amyloid-β Aggregates. J. Phys. Chem. B 2023, 127 (51), 10931–10940. 10.1021/acs.jpcb.3c06051.

(35) Mompeán, M.; Romano, V.; Pantoja-Uceda, D.; Stuani, C.; Baralle, F. E.; Buratti, E.; Laurents, D. V. The TDP-43 N-terminal Domain Structure at High Resolution. FEBS J. 2016, 283 (7), 1242–1260. 10.1111/febs.13651.

(36) Afroz, T.; Hock, E.-M.; Ernst, P.; Foglieni, C.; Jambeau, M.; Gilhespy, L. A. B.; Laferriere, F.; Maniecka, Z.; Plückthun, A.; Mittl, P.; Paganetti, P.; Allain, F. H. T.; Polymenidou, M. Functional and Dynamic Polymerization of the ALS-Linked Protein TDP-43 Antagonizes Its Pathologic Aggregation. Nat. Commun. 2017, 8 (1), 45. 10.1038/s41467-017-00062-0.

(37) Wright, G. S. A.; Watanabe, T. F.; Amporndanai, K.; Plotkin, S. S.; Cashman, N. R.; Antonyuk, S. V.; Hasnain, S. S. Purification and Structural Characterization of Aggregation-Prone Human TDP-43 Involved in Neurodegenerative Diseases. iScience 2020, 23 (6), 101159. 10.1016/j.isci.2020.101159.

(38) Sterling, T.; Irwin, J. J. ZINC 15 – Ligand Discovery for Everyone. J. Chem. Inf. Model. 2015, 55 (11), 2324–2337. 10.1021/acs.jcim.5b00559.

(39) Pınar, O.; Jana, A. K.; Yaşar, F. Stability and Dynamics of the N-Terminal Domain of TDP-43 and the Effect of Point Mutations. ACS Omega 2026, 11 (12), 19124–19133. 10.1021/acsomega.5c11735.

(40) Wang, H.; Logan, D. T.; Danielsson, J.; Oliveberg, M. Exposing the Distinctive Modular Behavior of β-Strands and α-Helices in Folded Proteins. Proc. Natl. Acad. Sci. 2020, 117 (46), 28775–28783. 10.1073/pnas.1920455117.

(41) Silberstein, S. D.; McCrory, D. C. Ergotamine and Dihydroergotamine: History, Pharmacology, and Efficacy. Headache J. Head Face Pain 2003, 43 (2), 144–166. 10.1046/j.1526-4610.2003.03034.x.

(42) Xu, Y.; Villalona-Calero, M. A. Irinotecan: Mechanisms of Tumor Resistance and Novel Strategies for Modulating Its Activity. Ann. Oncol. 2002, 13 (12), 1841–1851. 10.1093/annonc/mdf337.

(43) Ding, H. X.; Leverett, C. A.; Kyne, R. E.; Liu, K. K.-C.; Fink, S. J.; Flick, A. C.; O’Donnell, C. J. Synthetic Approaches to the 2013 New Drugs. Bioorg. Med. Chem. 2015, 23 (9), 1895–1922. 10.1016/j.bmc.2015.02.056.

(44) Oubrie, A.; Kaptein, A.; de Zwart, E.; Hoogenboom, N.; Goorden, R.; van de Kar, B.; van Hoek, M.; de Kimpe, V.; van der Heijden, R.; Borsboom, J.; Kazemier, B.; de Roos, J.; Scheffers, M.; Lommerse, J.; Schultz-Fademrecht, C.; Barf, T. Novel ATP Competitive MK2 Inhibitors with Potent Biochemical and Cell-Based Activity throughout the Series. Bioorg. Med. Chem. Lett. 2012, 22 (1), 613–618. 10.1016/j.bmcl.2011.10.071.

(45) Arkin, M. R.; Tang, Y.; Wells, J. A. Small-Molecule Inhibitors of Protein-Protein Interactions: Progressing towards the Reality. Chem. Biol. 2014, 21 (9), 1102–1114. 10.1016/j.chembiol.2014.09.001.

(46) Jo, S.; Kim, T.; Iyer, V. G.; Im, W. CHARMM-GUI: A Web-Based Graphical User Interface for CHARMM. J. Comput. Chem. 2008, 29 (11), 1859–1865. 10.1002/jcc.20945.

(47) Vanommeslaeghe, K.; Hatcher, E.; Acharya, C.; Kundu, S.; Zhong, S.; Shim, J.; Darian, E.; Guvench, O.; Lopes, P.; Vorobyov, I.; Mackerell Jr., A. D. CHARMM General Force Field: A Force Field for Drug-like Molecules Compatible with the CHARMM All-Atom Additive Biological Force Fields. J. Comput. Chem. 2010, 31 (4), 671–690. 10.1002/jcc.21367.

(49) Vanommeslaeghe, K.; Raman, E. P.; MacKerell, A. D. Automation of the CHARMM General Force Field (CGenFF) II: Assignment of Bonded Parameters and Partial Atomic Charges. J. Chem. Inf. Model. 2012, 52 (12), 3155–3168. 10.1021/ci3003649.

(50) Lindahl; Abraham; Hess; Spoel, van der. GROMACS 2021.4 Manual. 2021. 10.5281/zenodo.5636522.

(51) Huang, J.; Rauscher, S.; Nawrocki, G.; Ran, T.; Feig, M.; de Groot, B. L.; Grubmüller, H.; MacKerell, A. D. CHARMM36m: An Improved Force Field for Folded and Intrinsically Disordered Proteins. Nat. Methods 2017, 14 (1), 71–73. 10.1038/nmeth.4067.

(52) Jorgensen, W. L.; Chandrasekhar, J.; Madura, J. D.; Impey, R. W.; Klein, M. L. Comparison of Simple Potential Functions for Simulating Liquid Water. J. Chem. Phys. 1983, 79 (2), 926–935. 10.1063/1.445869.

(53) Darden, T.; York, D.; Pedersen, L. Particle Mesh Ewald: An *N* ⋅log(*N*) Method for Ewald Sums in Large Systems. J. Chem. Phys. 1993, 98 (12), 10089–10092. 10.1063/1.464397.

(54) Berendsen, H. J. C.; Postma, J. P. M.; Van Gunsteren, W. F.; DiNola, A.; Haak, J. R. Molecular Dynamics with Coupling to an External Bath. J. Chem. Phys. 1984, 81 (8), 3684–3690. 10.1063/1.448118.

(55) Parrinello, M.; Rahman, A. Polymorphic Transitions in Single Crystals: A New Molecular Dynamics Method. J. Appl. Phys. 1981, 52 (12), 7182–7190. 10.1063/1.328693.

(56) Humphrey, W.; Dalke, A.; Schulten, K. VMD: Visual Molecular Dynamics. J. Mol. Graph. 1996, 14 (1), 33–38. 10.1016/0263-7855(96)00018-5.

(57) McGibbon, R. T.; Beauchamp, K. A.; Harrigan, M. P.; Klein, C.; Swails, J. M.; Hernández, C. X.; Schwantes, C. R.; Wang, L.-P.; Lane, T. J.; Pande, V. S. MDTraj: A Modern Open Library for the Analysis of Molecular Dynamics Trajectories. Biophys. J. 2015, 109 (8), 1528–1532. 10.1016/j.bpj.2015.08.015.

(58) Best, R. B.; Hummer, G.; Eaton, W. A. Native Contacts Determine Protein Folding Mechanisms in Atomistic Simulations. Proc. Natl. Acad. Sci. 2013, 110 (44), 17874–17879. 10.1073/pnas.1311599110.

(59) Scheurer, M.; Rodenkirch, P.; Siggel, M.; Bernardi, R. C.; Schulten, K.; Tajkhorshid, E.; Rudack, T. PyContact: Rapid, Customizable, and Visual Analysis of Noncovalent Interactions in MD Simulations. Biophys. J. 2018, 114 (3), 577–583. 10.1016/j.bpj.2017.12.003.

(60) Valdés-Tresanco, M. S.; Valdés-Tresanco, M. E.; Valiente, P. A.; Moreno, E. gmx_MMPBSA: A New Tool to Perform End-State Free Energy Calculations with GROMACS. J. Chem. Theory Comput. 2021, 17 (10), 6281–6291. 10.1021/acs.jctc.1c00645.

(61) Schrödinger, LLC. The PyMOL Molecular Graphics System, Version 1.8, 2015.

(62) Zhou, X.; Zheng, W.; Li, Y.; Pearce, R.; Zhang, C.; Bell, E. W.; Zhang, G.; Zhang, Y. I-TASSER-MTD: A Deep-Learning-Based Platform for Multi-Domain Protein Structure and Function Prediction. Nat. Protoc. 2022, 17 (10), 2326–2353. 10.1038/s41596-022-00728-0.

(63) O’Boyle, N. M.; Banck, M.; James, C. A.; Morley, C.; Vandermeersch, T.; Hutchison, G. R. Open Babel: An Open Chemical Toolbox. J. Cheminformatics 2011, 3 (1), 33. 10.1186/1758-2946-3-33.

(64) Trott, O.; Olson, A. J. AutoDock Vina: Improving the Speed and Accuracy of Docking with a New Scoring Function, Efficient Optimization, and Multithreading. J. Comput. Chem. 2010, 31 (2), 455–461. 10.1002/jcc.21334.

(65) Morris, G. M.; Huey, R.; Lindstrom, W.; Sanner, M. F.; Belew, R. K.; Goodsell, D. S.; Olson, A. J. AutoDock4 and AutoDockTools4: Automated Docking with Selective Receptor Flexibility. J. Comput. Chem. 2009, 30 (16), 2785–2791. 10.1002/jcc.21256.

