## Supplementary Information for "Mechanistic Insights into Unfolding of the Mitochondrial Localization Sequence M1 of TDP-43 and Identification of a MAPKAPK2 Inhibitor as a potential binder of M1 for inhibition of TDP-43’s Aberrant Mitochondrial Localization"

Title

Basant K. Patel, PhD

* Corresponding author

Himanshu Joshi, PhD

### Supplementary Figures

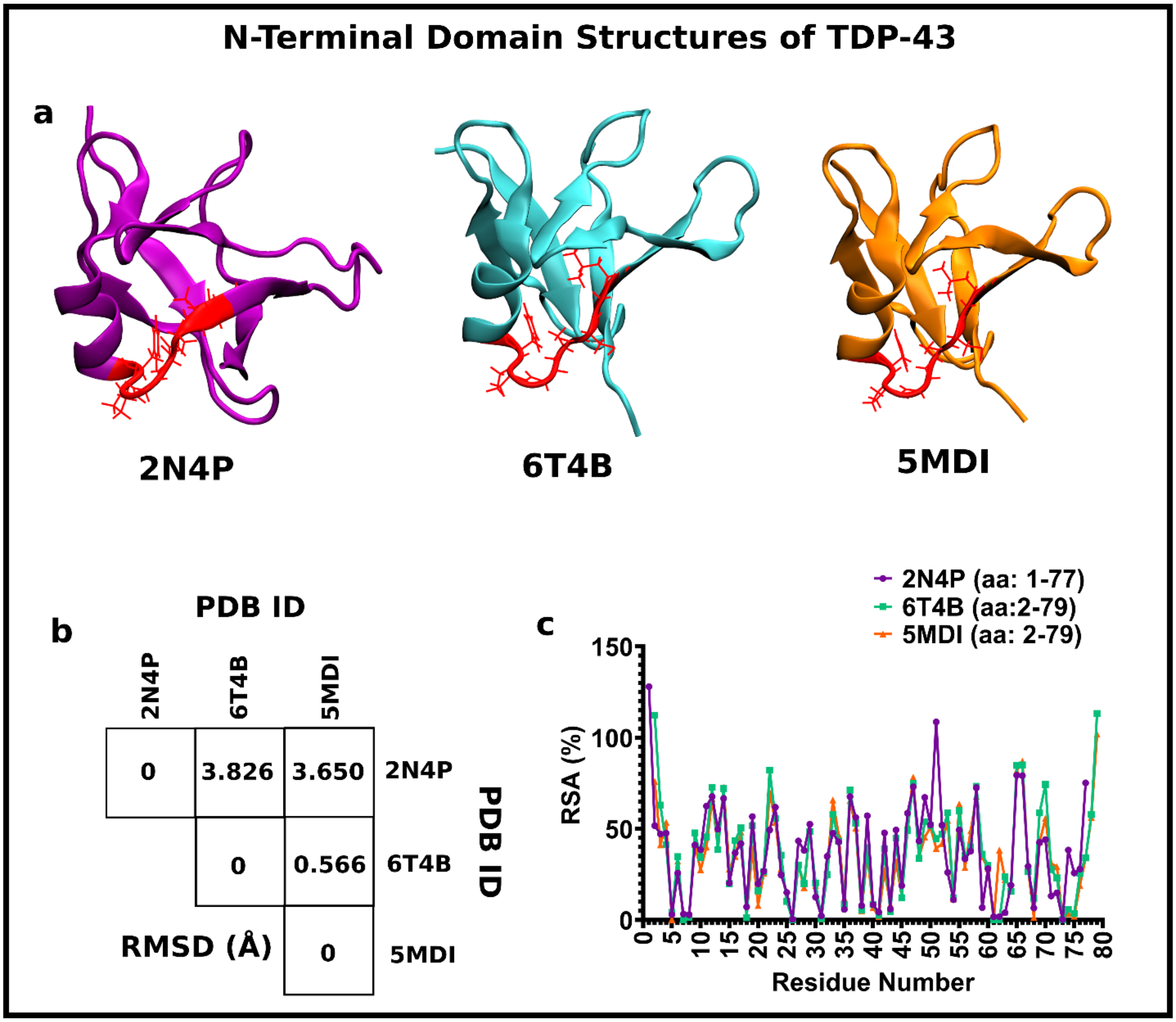


**Figure SF1 - The comparison of the N-terminal domain structures of TDP-43 used in the current study.**

a) The structures of the N-terminal domain (NTD) of TDP-43 elucidated by solution NMR (PDB ID: 2N4P; aa: 1-77), X-ray crystallography (chain A of PDB ID: 5MDI, aa: 2-79 and chain A of PDB ID: 6T4B: 2-79). The internal mitochondrial signal sequence, M1, is shown in red as a stick, along with the new cartoon representation for the remaining region. b) The matrix of the RMSD comparison among the different N-terminal structures used. The C-α RMSD for the atoms of residue 2-77 for all the structures after aligning are only considered. The matrix shows that the backbone C-α RMSD for the two X-ray crystallography structures (PDB ID: 5MDI, A-chain; 6T4B, A-chain) doesn't vary much among themselves but they vary significantly with the solution NMR structure (PDB ID: 2N4P) indicative of the structural difference. However, no difference in the overall fold of the protein regions is visible among these structures. c) The Percentage Relative Solvent Accessibility (%RSA) value for the residues in the different N-terminal structures used. Most residues have comparable %RSA, confirming the similar folds among the domain structures.

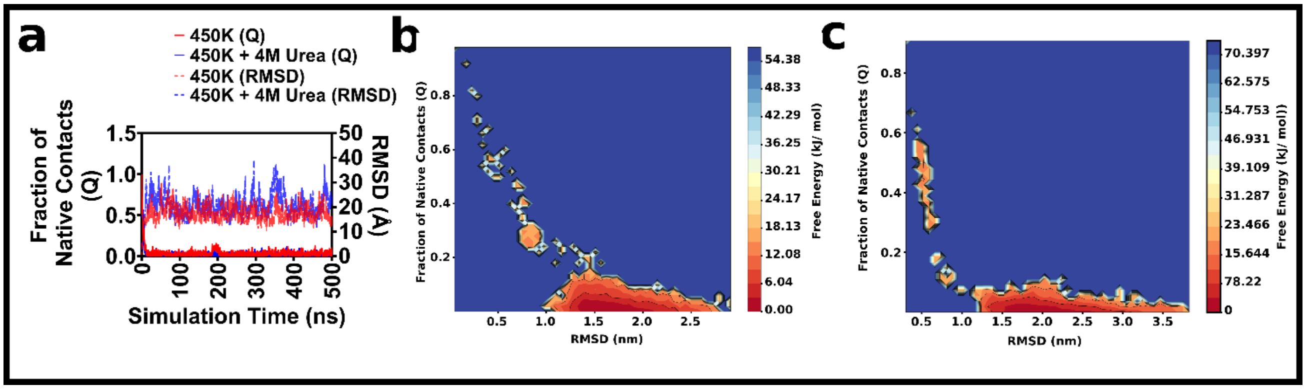


**Figure SF2 - MD simulation analysis of the solution NMR structure of the N-terminal domain (NTD) of TDP-43 (PDB ID: 2N4P, aa: 1-77) at 450 K and 450 K + 4 M urea simulation conditions**: a) The fraction of native contacts (Q) and RMSD profile of the 2N4P at 450 K and 450 K + 4 M urea conditions. The Q and RMSD profile indicate complete instantaneous unfolding of the protein in these two conditions b) Free energy contour plot calculated as a function of the joint probability of Q and RMSD at 450 K. c) Free energy contour plot calculated as a function of the joint probability of Q and RMSD at 450 K + 4 M urea; The free energy change, $\Delta G = -RT \ln P(Q, RMSD)$, where R is Universal gas constant (8.314 J mol-1 K-1), T is temperature in K, and P(Q, RMSD) is the joint probability of Q and RMSD. The free energy change is normalized to the minimum value.

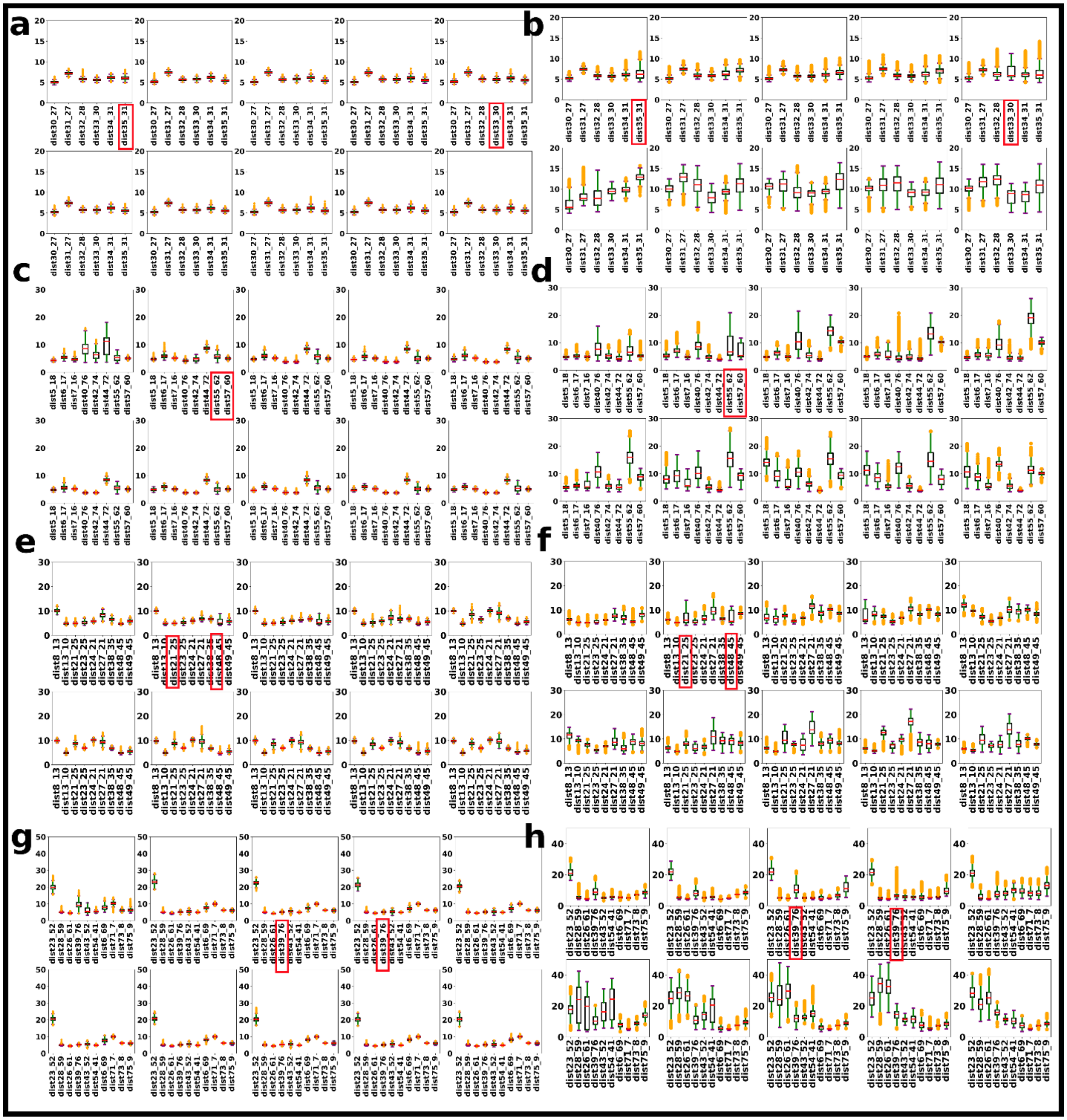


**Figure SF3: Pair distances between the residues in hydrogen bonding throughout the simulation time for the solution NMR structure of the N-terminal domain (NTD) of TDP-43 (PDB ID: 2N4P).**

a-b) The pair distances between the residues in the α-helix region of the N-terminal domain (NTD) structure of TDP-43 (PDB ID: 2N4P) reported to form hydrogen bonding at 310 K and 350 K + 4

M, respectively. The median, interquartile ranges, and standard deviations are represented by the middle line, the box, and the whiskers, respectively, with outliers shown as solid circles in the box plot. These values are computed every 50 ns and displayed in the panels. c-d) The pair-distance distribution for the residues from the beta-sheets that form the beta bridges in the 2N4P structure at 310 K and 350 K + 4 M, respectively. e-f) The pair-distance distribution for the residues in the loop region of the 2N4P structure at 310 K and 350 K + 4 M, respectively. g-h) Pair-distance distribution for the residues that are far away in the N-terminal sequence but are reported to form hydrogen bonding at 310 K and 350 K + 4 M, respectively. The residue pairs in the orange boxes corresponds to those which show increased inter-quartile range (IQR) or increased outliers at 350 K + 4 M urea compared to the 310 K suggesting the affected hydrogen-bonding. The residue-pairs involving Phe-35, and those between β4-β5 in hydrogen-bonding were affected at early simulation time indicating their role in early unfolding.

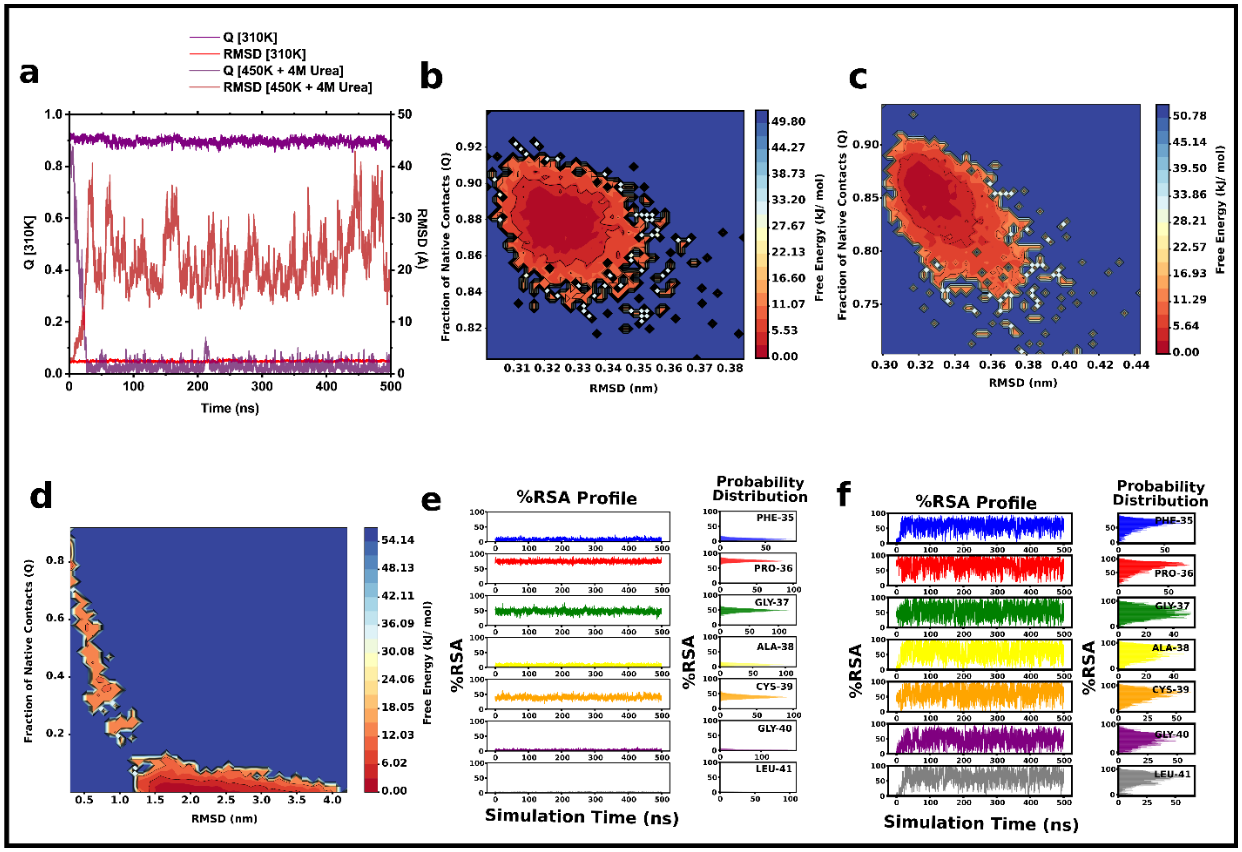


**Figure SF4 -MD simulation analysis of the X-ray crystallographic structure of the N-terminal domain (NTD) of TDP-43 (PDB ID: 6T4B) at 310 K, 350 K, 400 K + 4 M urea, and 450 K + 4 M urea simulation conditions**.

a) The fraction of native contacts (Q) and RMSD profile of the 6T4B at 310 K and 450 K + 4 M urea conditions. The Q and RMSD profile suggest the instantaneous unfolding of the protein structures at 450 K + 4 M urea. b-d) The free energy profile based on the joint probability of Q and RMSD for 6T4B at 350 K, 400K + 4M, and 450K + 4M urea conditions, respectively; The free energy change, ΔG = -RT ln P(Q, RMSD), where R is Universal gas constant (8.314 J mol-1 K-1), T is temperature in K, and P(Q, RMSD) is the joint probability of Q and RMSD. The free energy change is normalized to the minimum value. The free energy profiles at 350 K, and 400 K + 4 M urea show that the protein remains folded in the two simulation conditions. However, the free energy profiles at 450 K + 4 M urea indicates complete unfolding instantaneously without any free energy basin corresponding to the intermediate states. e) Percentage relative solvent accessibility of the residues of the internal mitochondrial region, M1 at 310 K. f) Percentage relative solvent accessibility of the residues of the M1 region in the 450 K + 4 M urea. The %RSA at 450 K + 4 M urea of the M1 indicates complete solvent exposure of the region at 450 K + 4 M urea.

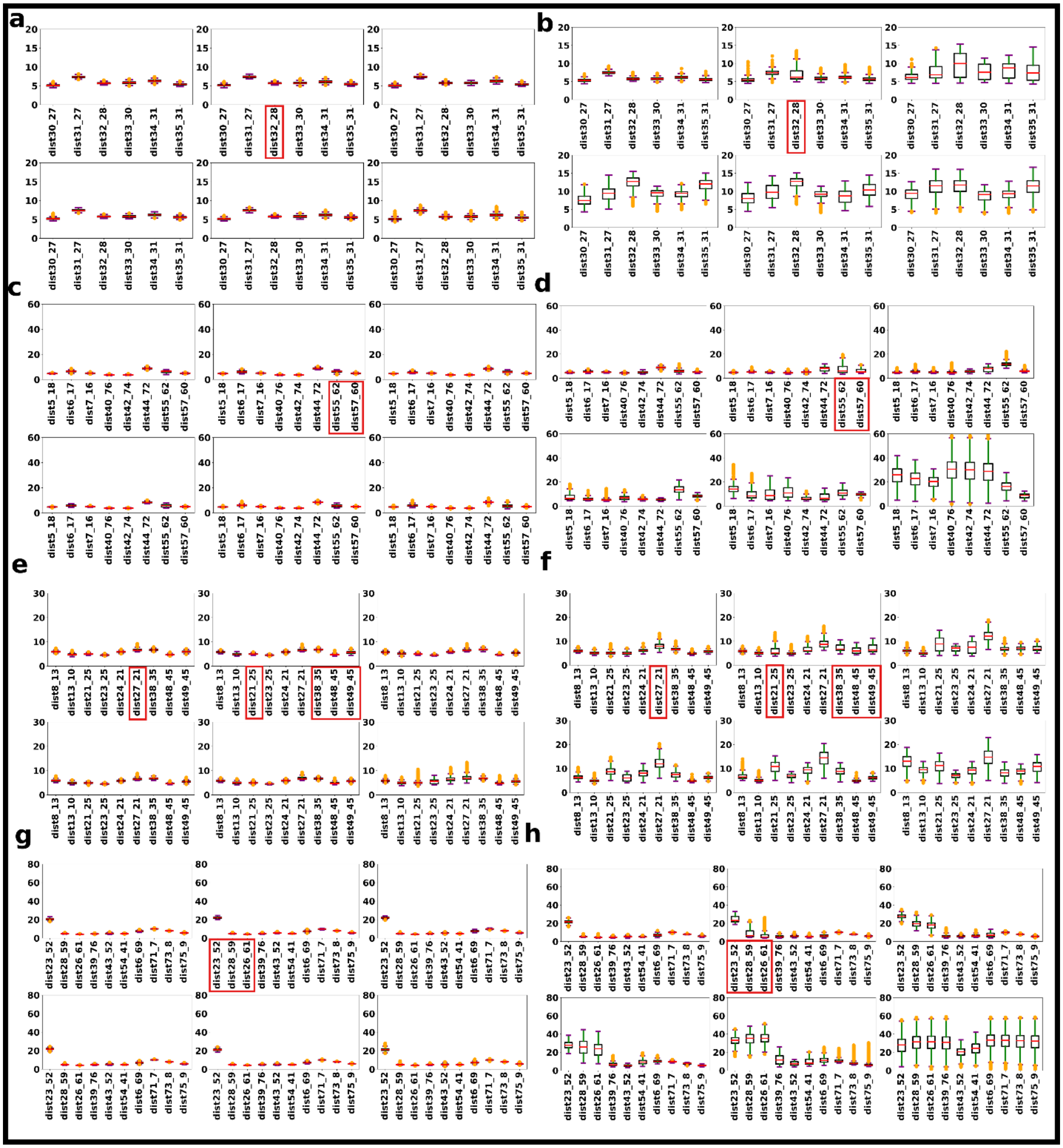


**Figure SF5- Pair distances between the residues in hydrogen bonding throughout the simulation time for the X-ray crystallographic structure of N-terminal domain (NTD) of TDP-43 (PDB ID: 6T4B).**

a-b) The pair distances between the residues in the α-helix region of the 6T4B structure reported to form hydrogen bonding at 310 K and 450 K+4 M, respectively. The median, interquartile ranges, and standard deviations are represented by the middle line, the box, and the whiskers, respectively, with outliers shown as solid circles in the box plot. These values are computed every 5 ns until 30 ns and are shown as panels. 30 ns is considered as the structure completely unfolded at this simulation time point c-d) The pair-distance distribution for the residues from the beta-sheets that form the beta bridges in the 6T4B structure at 310 K and 450 K+4 M, respectively. e-f) The pair-distance distribution for the residues in the loop region of the 6T4B structure at 310 K and 450 K+4 M, respectively. g-h) Pair-distance distribution for the residues that are far away in the N-terminal sequence but are reported to form hydrogen bonding at 310 K and 450 K+4 M, respectively. The residue pairs in the orange boxes corresponds to those which show increased inter-quartile range (IQR) or increased outliers at 350 K + 4 M urea compared to the 310 K suggesting the affected hydrogen-bonding. The residue-pairs involving Phe-35, and those between β4-β5 in hydrogen-bonding were affected at early simulation time indicating their role in early unfolding.

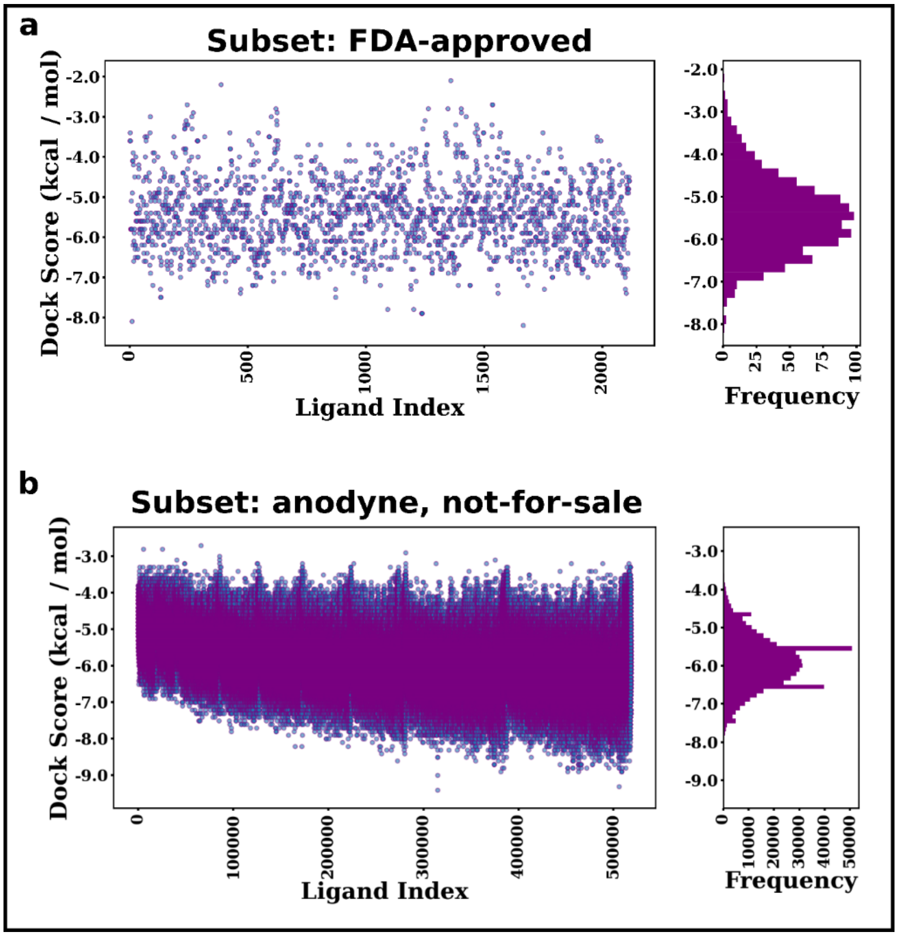


**Figure SF6 - Virtual screening of FDA-approved/non-FDA-approved drugs from an online small molecule library (ZINC15).**

a) Scatter plot of the defined docking scores for the FDA-approved small molecules against the internal mitochondrial region, M1 (aa: 35 – 41) in the crystal structure of the N-terminal domain (NTD) of TDP-43 (PDB ID: 5MDI). b) The scatter plot for the non-FDA-approved small molecules under the category annotated, not-for-sale, against the M1 region in the N-terminal domain of TDP-43.

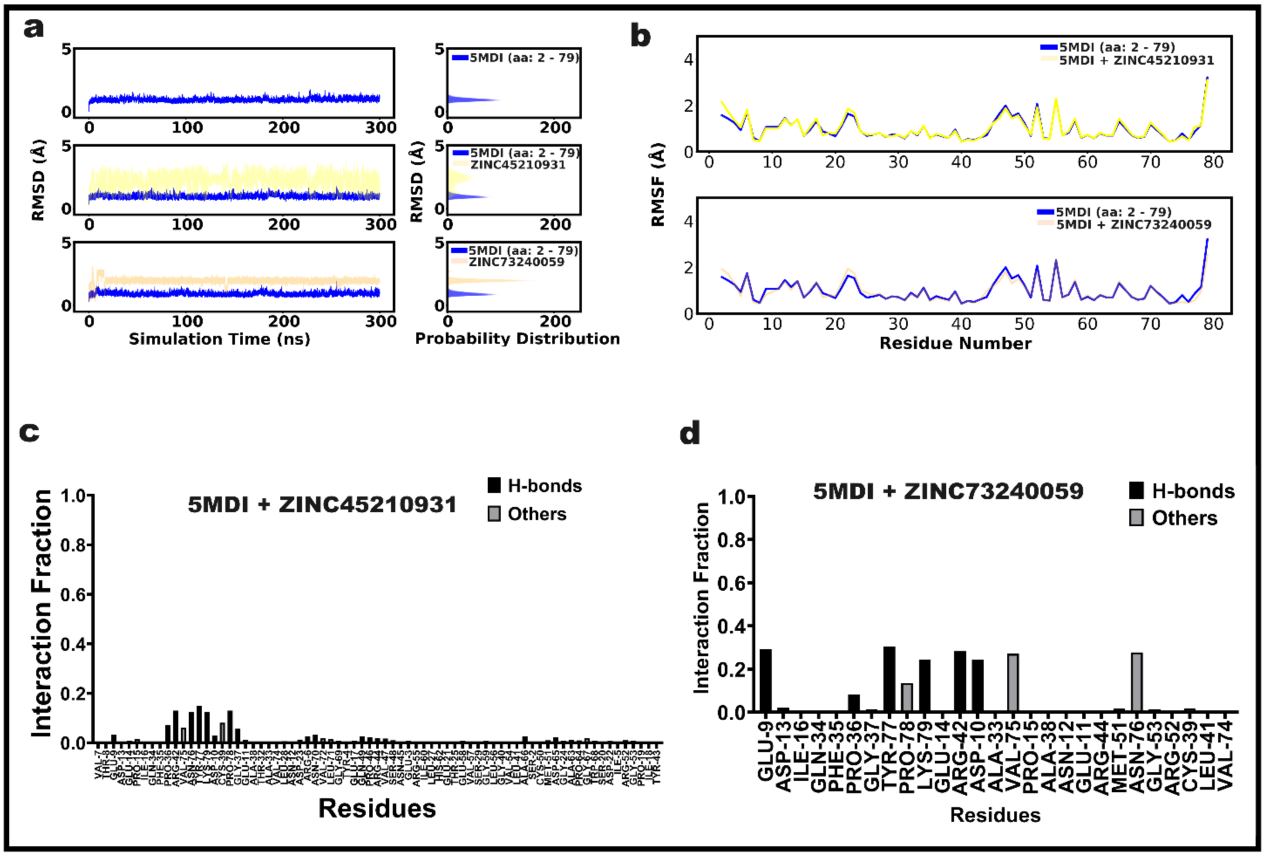


**Figure SF7- 300 K simulation results for the x-ray crystallographic structure (5MDI) of the N-terminal of TDP-43 with top-ranked molecules from the virtual screening.**

a) RMSD profile of the protein-alone and protein-ligand simulations performed on the top molecules from the virtual screening. b) RMSF profile of the residues in protein-ligand compared against the protein-alone simulations. c-d) Interaction profiles for the 5MDI with the small molecules ZINC45210931 and ZINC73240059 from the annotated class of molecules virtually screened. The molecule ZINC73240059, a putative MAPAPK2 inhibitor, interacted with the residues of M1 (aa: 35-41) in most of the frames of the simulation.

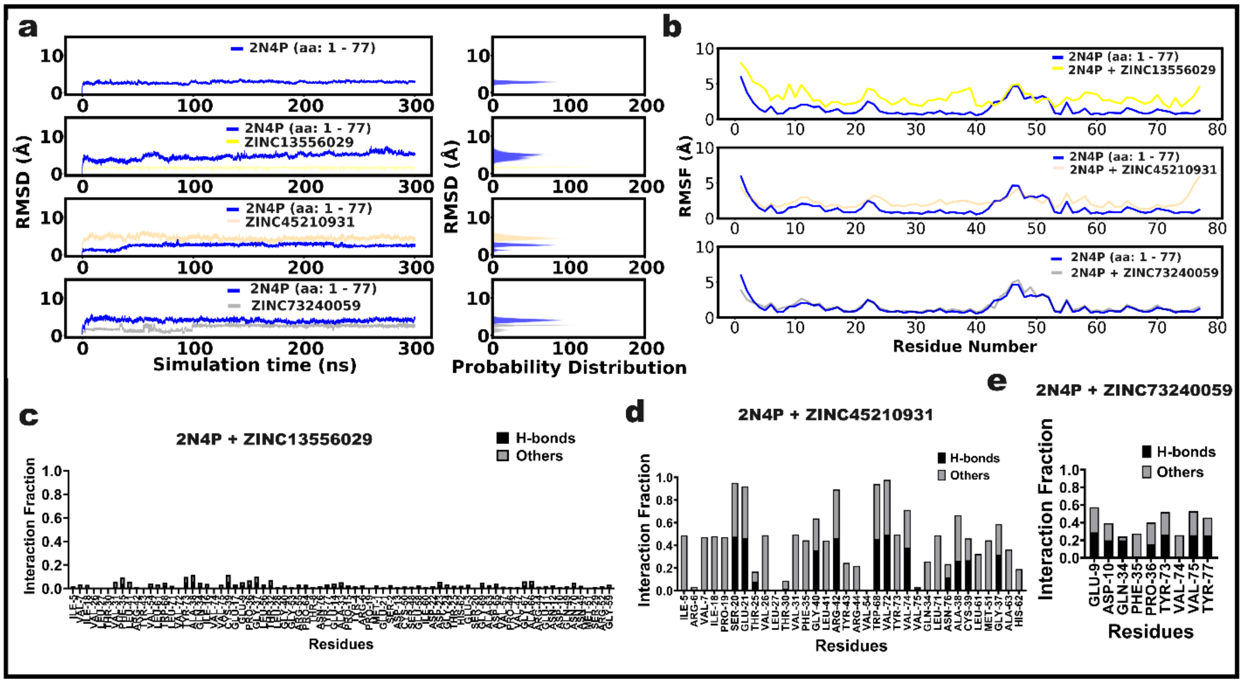


**Figure SF8- 300 K simulation results for the solution NMR structure (PDB ID: 2N4P) of the N-terminal domain (NTD) of TDP-43 with top-ranked molecules from the virtual screening.**

a) RMSD profile of the protein-alone and protein-ligand simulations performed on the top molecules from the virtual screening. b) RMSF profile of the residues in protein-ligand compared against the protein-alone simulations. c-d) Interaction profiles for the 2N4P with the small molecules ZINC13556029, ZINC45210931, and ZINC73240059 from the non-FDA-approved molecules labeled as annotated class of molecules in the ZINC15 library that is virtually screened. ZINC13556029 is used as a negative control for the comparison. The molecule ZINC73240059, a putative MAPAPK2 inhibitor, interacted stably at M1 for most of the simulation time and showed little RMSF relative to the protein alone, compared to the other molecules.

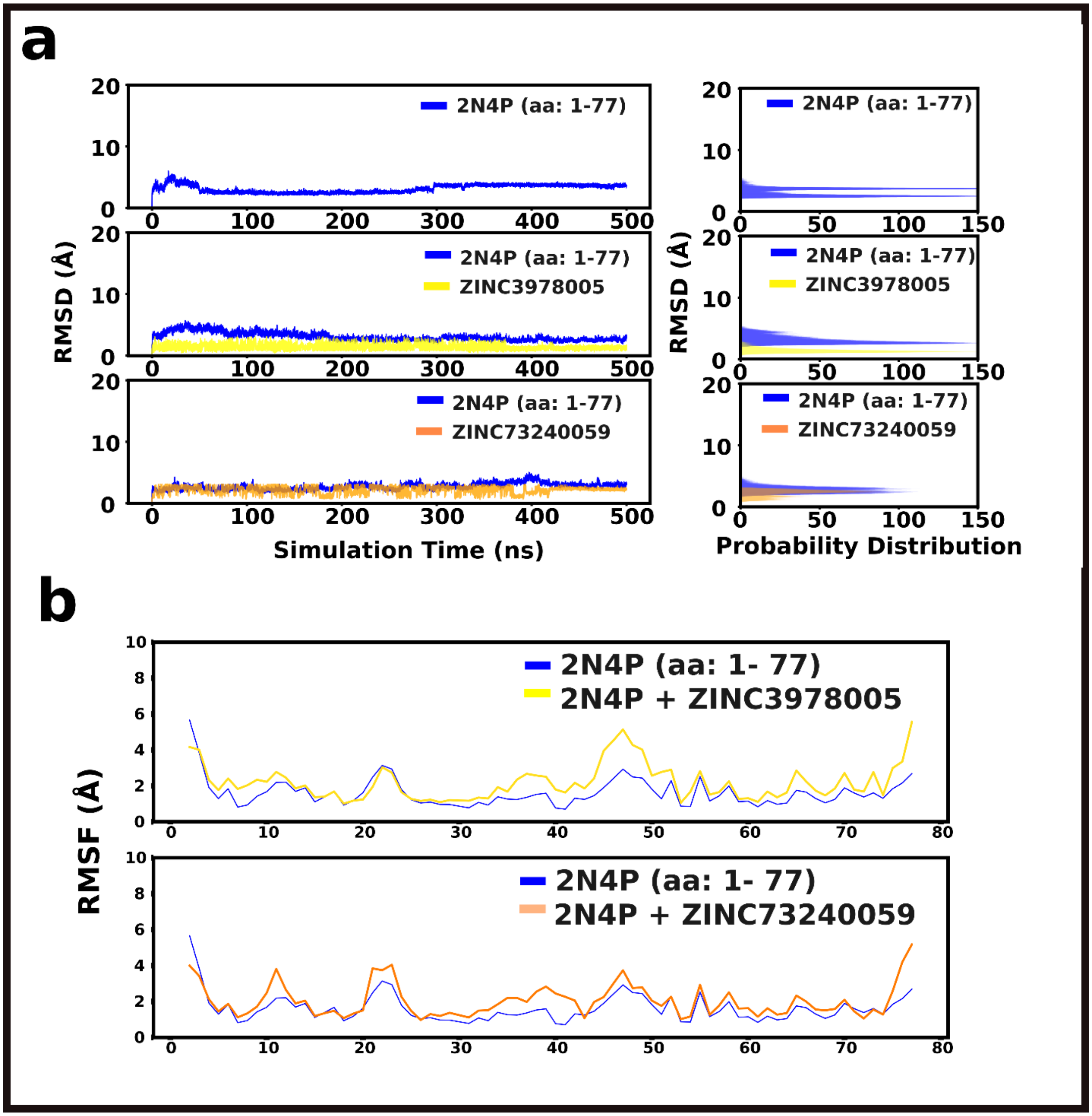


**Figure SF9 – RMSD, RMSF profiles of the complex of the solution NMR structure of the N-terminal domain (NTD) of TDP-43 (PDB ID: 2N4P) with the top two small molecules from screening studied at 310 K.**

a) RMSD profile of the small molecules ZINC3978005 (Dihydroergotamine) and ZINC73240059 studied for their interaction with the solution NMR structure of the N-terminal domain of TDP-43

(PDB ID: 2N4P) at 310K. b) RMSF profile of the two molecules studied with the 2N4P structure at 310K. The RMSD and RMSF profiles showed fewer fluctuations for the 2N4P in complex with ZINC73240059 than for the protein alone simulations, whereas ZINC3978005 showed increased fluctuations.

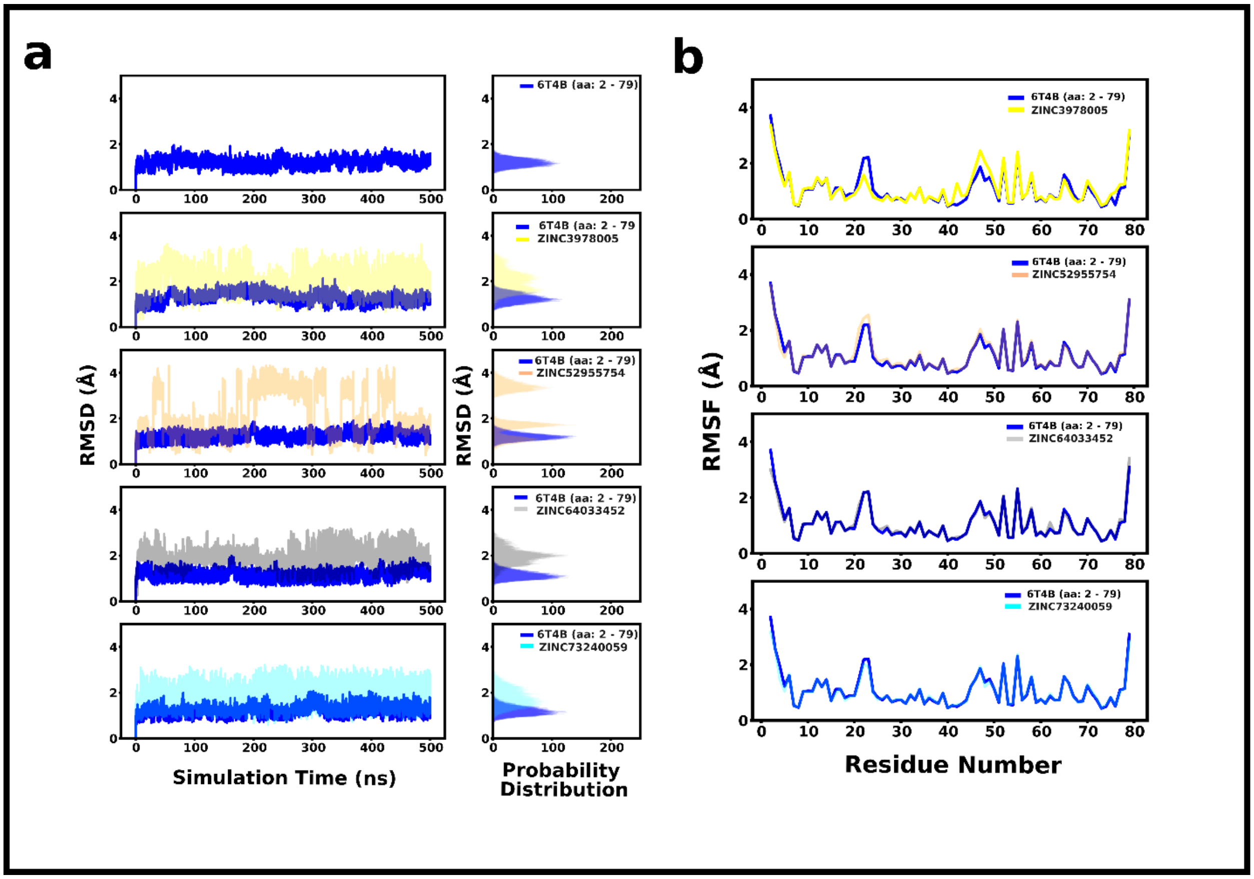


**Figure SF10 – RMSD, RMSF profiles of the complex of the X-ray crystallographic structure of the N-terminal domain (NTD) of TDP-43 (PDB ID: 6T4B) structure with small molecules from screening studied at 310K.**

a) RMSD profile of the small molecules from screening studied for their interaction with the X-ray crystallographic structure of the N-terminal domain (NTD) of TDP-43 (PDB ID: 6T4B) at 310K. b) RMSF profile of the complex of the studied molecules with the 6T4B structure at 310K. The RMSD and RMSF profiles showed fewer fluctuations for the 2N4P in complex with ZINC73240059 than for the protein alone simulations.

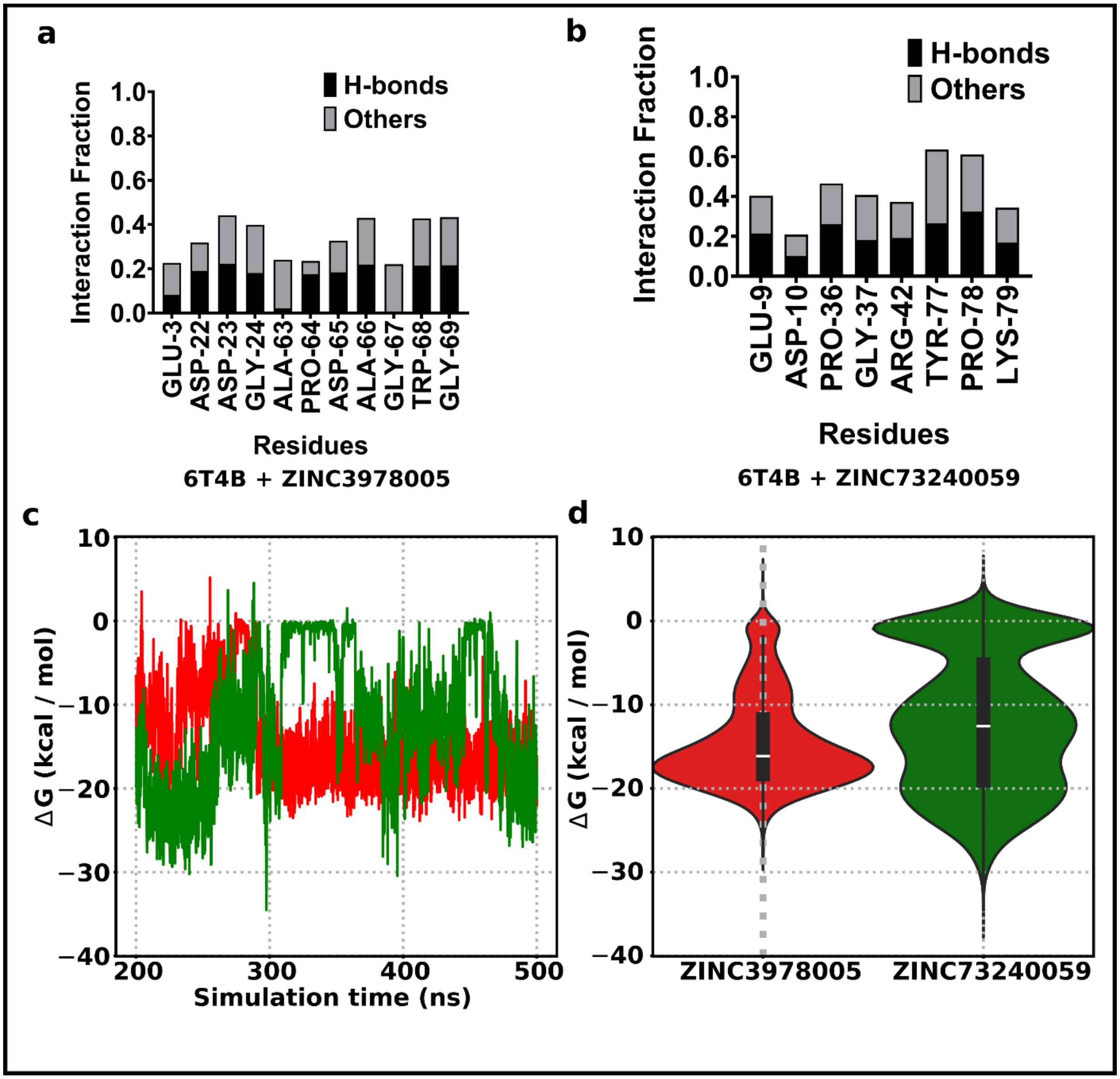


**Figure SF11 – MD simulation analysis of the virtually screened molecules with the crystallographic structure of the N-terminal domain (NTD) of TDP-43 (PDB ID: 6T4B).**

a-b) The interaction fraction profile of the 6T4B with the small molecules ZINC3978005 (Dihydroergotamine), ZINC52955754 (Ergotamine), and ZINC7340059. The interaction fraction of 1 means the residue of the protein interacts with the small molecule with both the hydrogen bond and other non-covalent interactions throughout the simulation time (H-bond + Other bonds

= 0.5 + 0.5 = 1.0). c-d) Free energy profile for the protein (6T4B) with ZINC3978005 and ZINC73240059 shown as a line plot in the left panel and the probability distribution of the values as a violin plot in the right panel. The interaction profile suggests a stable interaction of the M1 residues with the molecule ZINC73240059, resulting in the lowest negative free energy compared to Dihydroergotamine and Ergotamine.

### Supplementary Tables

**Table S1:** The docking parameters used for the defined docking (virtual screening) against the internal mitochondrial sequence, M1, in the N-terminal domain (NTD) of TDP-43, and the blind docking experiments.

| **Parameters** | **Virtual Screening (Defined Docking against M1)** **AutoDock Vina v1.12** | **Blind Docking** **AutoDock 4.2.6** | | | | | | | | | |
|---|---|---|---|---|---|---|---|---|---|---|---|
| **Receptor Macromolecule** | **5MDI** | **Modelled FL-TDP-43#** | **5MDI** | **2N4P** | **Cluster Representative Structures*** | | | | | | |
| | | | | | **0–50 ns** | **50 – 100 ns** | **100 – 150 ns** | **150 – 200 ns** | **200 – 250 ns** | **250 – 300 ns** | **300 – 500 ns** |
| **Size_x** | 40 | 96 | 104 | 126 | 108 | 114 | 108 | 108 | 114 | 96 | 96 |
| **Size_y** | 62 | 126 | 96 | 96 | 86 | 82 | 86 | 86 | 82 | 102 | 102 |
| **Size_z** | 46 | 126 | 88 | 94 | 104 | 102 | 104 | 104 | 102 | 112 | 112 |
| **Center_x** | -1.472 | -11.296 | -5.517 | -5.592 | 0.501 | 0.478 | 0.57 | 0.57 | 0.481 | 0.177 | 0.176 |
| **Center_y** | -32.636 | -57.245 | -34.81 | -34.81 | 0.106 | 0.177 | -0.077 | -0.077 | 0.096 | -0.035 | 0.007 |
| **Center_z** | -7.381 | 33.665 | 5.592 | -5.592 | -0.134 | -0.071 | -0.102 | -0.102 | -0.163 | -0.294 | -0.242 |
| **Exhaustiveness** | 8 | NA | NA | | | | | | | | |
| **Dielectric constant** | NA | -0.1465 | -0.1465 | | | | | | | | |
| **No. of GA runs** | NA | 50 | 10 | | | | | | | | |

*Protein structure alone after clustering, taken from different simulation time intervals in the 350 K + 4M Urea simulations of 2N4P.

#Full-length TDP-43 modeled in the I-TASSER MTD webserver, and the grid spacing for the simulation is set to 0.8.

**Table S2:** The RMSD of the modeled full-length TDP-43 with the solved domain structures of TDP-43 from RCSB-PDB as reference structures.

| **Domain Structures of TDP-43** | **FL-TDP-43*** |
|---|---|
| 2N4P<br>N-terminal<br>(aa: 1-77) | 6.1 |
| 6T4B<br>N-terminal<br>(aa: 2-79) | 7.23 |
| 4BS2<br>RRM1-2<br>(aa: 102-269) | 11.02 |

* RMSD for Modeled Full-length TDP-43 from I-TASSER-MTD with reference to the different domain structures available in PDB (Å)

**Table S3:** Docking scores for the putative MAPAPK2 inhibitor, ZINC73240059, bound to the internal mitochondrial sequence, M1, in the cluster representative structures from unfolding simulations of the N-terminal domain (NTD) of TDP-43 (PDB ID: 2N4P).

| **Cluster Representative Structures interval*** | **AutoDock 4.2 score (kcal/mol)#** | **Inhibitory constant $K_i$ (μM)** | **Residues of M1 involved and the type of interaction** |
|---|---|---|---|
| 0 – 50 ns | -8.22 | 0.94 | Van der Waals: Phe-35 |
| 50 – 100 ns | -7.87 | 1.70 | Van der Waals: Phe-35, Pro-36, Gly-37, Ala-38, Cys-39 |
| 150 – 200 ns | -7.14 | 5.85 | Van der Waals: Phe-35, Pro-36 |
| 200 – 250 ns | -7.12 | 4.28 | Van der Waals: Gly-40, Leu-41 |
| 250 – 300 ns | -7.81 | 1.89 | Van der Waals: Leu-41 |
| 300 – 500 ns | -5.99 | 40.44 | Van der Waals: Gly-37, Ala-38 |

* The time interval in which the cluster representative structures, from the unfolding simulation of the N-terminal domain structure (PDB ID: 2N4P) that were carried out at 350 K + 4 M Urea, were chosen.

### Lowest AutoDock 4.2 scores that were reported for the complex of ZINC73240059 interacting with the M1 residues (aa: 35-41) of the N-terminal domain of TDP-43.

**Table S4:** The components of the free energy calculations and their contributions as calculated by gmx_MMPBSA for the solution NMR structure of the N-terminal domain (NTD) of TDP-43 (PDB ID: 2N4P) with the molecules that stably interacted with the internal mitochondrial sequence, M1, in the MD simulations.

| **Energy Components** | **2N4P (aa: 1 – 77)** | | **6T4B (aa: 2 – 79)** | |
|---|---|---|---|---|
| | **ZINC3978005¶** | **ZINC73240059¶** | **ZINC3978005¶** | **ZINC73240059¶** |
| VDWAALS ($E_{VdW}$) | -17.26 ± 0.24 | -20.87 ± 0.27 | -20.66 ± 0.12 | -19.33 ± 0.13 |
| EEL ($E_{ele}$) | -77.5 ± 0.75 | -214.12 ± 1.85 | -111.75 ± 0.49 | -183.06 ± 1.08 |
| EPB ($E_{polar}$) | 85.09 ± 0.72 | 218.71 ± 1.87 | 120.44 ± 0.52 | 190.48 ± 1.08 |
| ENPOLAR ($E_{nonpolar}$) | -1.96 ± 0.03 | -2.49 ± 0.03 | -2.54 ± 0.01 | -2.33 ± 0.01 |
| GGAS ($E_{VdW} + E_{ele}$) | -94.75 ± 0.71 | -234.99 ± 2.07 | -132.41 ± 0.58 | -202.39 ± 1.15 |
| GSOLV ($E_{polar} + E_{nonpolar}$) | 83.13 ± 0.72 | 216.22 ± 1.85 | 117.9 ± 0.51 | 188.15 ± 1.07 |
| **Total ($E_{VdW} + E_{ele} + E_{polar} + E_{nonpolar}$)** | **-11.62 ± 0.16** | **-18.76 ± 0.25** | **-14.51 ± 0.1** | **-14.24 ± 0.1** |

¶The values are in kcal/mol ± Standard error of the means (SEM) over the trajectory frames used.

**Table S5:** Per residue decomposition to the overall free energy change for the X-ray crystallographic structure of the N-terminal domain of TDP-43 (PDB ID: 6T4B) with ZINC73240059 as calculated in gmx_MMPBSA.

| Residues | 6T4B + ZINC73240059* |
|---|---|
| PRO-36 | -0.65 ± 0.01 |
| TYR-77 | -2.38 ± 0.02 |
| PRO-78 | -0.9 ± 0.01 |
| ZINC73240059 | -10.1 ± 0.88 |

** ΔG values in kcal/mol ± Standard error values*

### Supplementary Movies

**Movie SM1 – MD simulations on the solution NMR structure of the N-terminal domain (NTD) of TDP-43; PDB ID: 2N4P (aa: 1-77) at different temperatures and denaturant conditions:** Protein alone simulations of the NMR structure of the N-terminal of TDP-43 (PDB ID: 2N4P; aa:1-77) at 310K, 350K + 4M urea, 450K, and 450K + 4M urea with the protein represented in cyan color and new cartoon representations for 310K, 450K, 450K + 4M urea. The protein is shown in a new cartoon representation for 350K + 4M urea, in purple. The internal mitochondrial sequence, the M1 region (aa 35-41), in all structures is colored grey.

**Link for the movie SM1:**

https://www.dropbox.com/scl/fi/g7xgyzl87yj4j5d7xkz3a/2N4P_Protein_alone_MDsimulation.mpeg?rlkey=0sdl5akml5ozk218vhq89jf5j&st=qkx5007r&dl=0

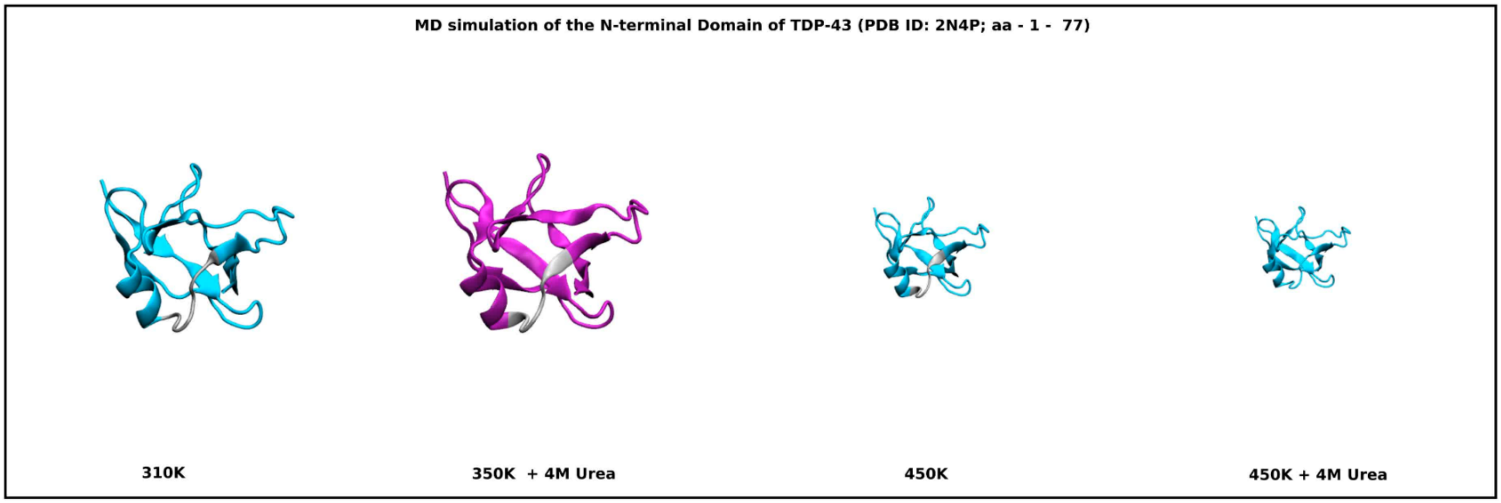

**Movie SM2 – MD simulations of the N-terminal domain structures of TDP-43 with screened small molecules at 310K:** Complex simulations of the NMR structure of the N-terminal of TDP-43 (PDB ID: 2N4P; aa:1-77; 6T4B, aa – 2-79) at 310K with the protein represented in purple color and new cartoon representations. The M1 region (aa 35-41) is colored yellow. The molecules ZINC3978005 and ZINC73240059 were shown in VDW representation. The simulations show that ZINC73240059 stably interacts with the different N-terminal structures at M1 in most frames, whereas ZINC3978005 does not remain bound to M1 in most frames.

**Link for the movie SM2:**

https://www.dropbox.com/scl/fi/zbsxkdxw4t1yfeqt78a52/NTerm_SmallMolecules.mpeg?rlkey=6ledod6jb5quj11ef5fdq5j2d&st=wrnzfs1r&dl=0

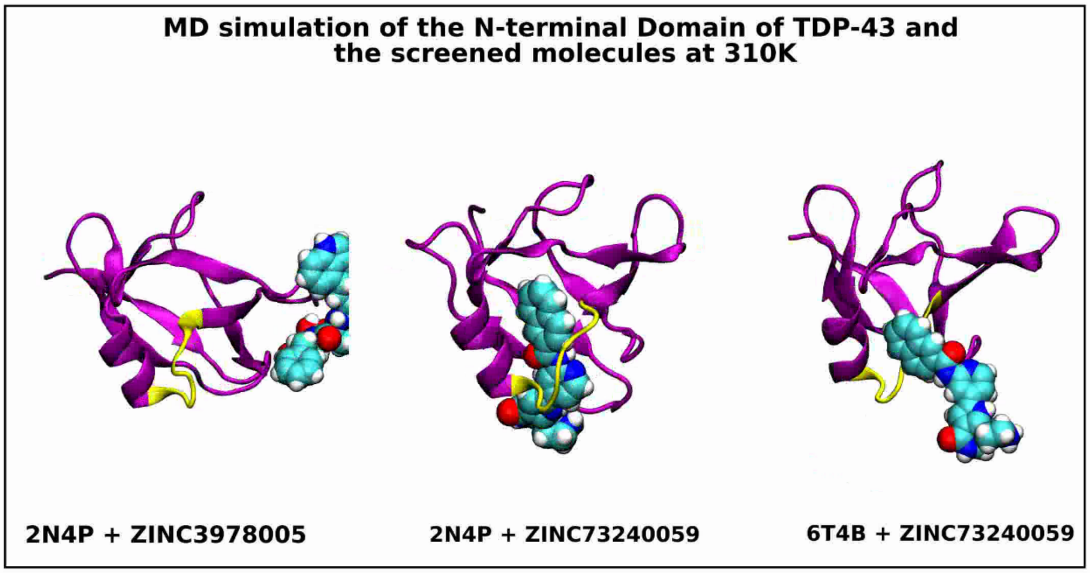
